# Plasmids link antibiotic resistance genes and phage defense systems in *E. coli*

**DOI:** 10.1101/2025.07.25.666796

**Authors:** Weronika Ślesak, Piotr Jedryszek, Daniel Cazares, William Matlock, R. Craig MacLean

## Abstract

Phage therapy has been proposed as an alternative to antibiotics to treat resistant infections. However, we have a limited understanding of how antibiotic resistance genes (ARGs) associate with bacterial phage defense systems (PDSs). Here, we explore the relationship between ARGs and PDSs in a sample of 2,559 plasmids originating from 1,044 *E. coli* isolates, representing a snapshot of clinical and non-clinical diversity in Oxfordshire, UK (2008-2020). In total, we identify 3,193 ARGs and 14,013 PDSs (180 unique types). We demonstrate that *E. coli* plasmids are enriched for ARGs and PDSs (both p<0.001), with a bias towards toxin-antitoxin/abortive-infection, TIR-domain and CBASS systems (all q<0.025). We proceed to show that ARGs and PDSs are physically linked by plasmids (*p*<0.001). Together, our results suggest that phage therapy may inadvertently select for antibiotic resistant bacteria, and that antibiotic use may similarly drive resistance to phage.

## Introduction

Bacterial defense systems can protect the cell against invaders, such as plasmids and phage^1^. Plasmids are extrachromosomal replicons often able to transfer between bacterial hosts via conjugation^2^. Phage, and in particular, lytic phage, are bacterial viruses that, like plasmids, rely on the host cell for replication, but unlike plasmids, ultimately kill the host during the infection cycle^3^. Plasmids are often associated with antimicrobial resistance genes (ARGs), which can confer resistance to the host cell, sometimes at a low fitness cost^4,5^. Moreover, plasmids can also encode phage defense systems (PDSs), such as toxin-antitoxin, restriction modification, and CRISPR-Cas systems, blurring the boundary between selfish genetic elements and mutualistic partners^6–9^.

One unresolved issue is the extent to which ARGs and PDSs co-occur on plasmids. This is important because the physical linkage of different resistance determinants can drive co-selection: selective pressure for one trait (phage resistance) may inadvertently maintain another (antibiotic resistance), and vice versa. Co-selection is already well documented for ARGs and other resistance genes, such as those against heavy metals or biocides, where a single stressor can select for multi-resistant bacteria^10,11^. Some examples have been reported of ARGs and PDSs co-localising on the same mobile genetic element, yet whether this occurs more generally remains to be seen^12–14^. Understanding these patterns is particularly relevant as phage therapy gains traction as a strategy to combat antibiotic resistant infections^15,16^.

In this study, we investigated the relationship between ARGs and PDSs in a large sample of 2,559 plasmids from 1,044 clinical and non-clinical *E. coli* isolates^17^. We first annotate the genomes for ARGs and PDSs, then explore (i) their overall enrichment on plasmids versus chromosomes, (ii) the enrichment of specific PDS types on plasmids versus chromosomes, and (iii) their linkage on plasmids.

## Methods

### Dataset curation

Isolates and associated genome assemblies were from an existing, pre-collated dataset^17^. The original dataset contained Enterobacterales species collected from within 60km in Oxfordshire from 2008-2020, covering multiple clinical and non-clinical niches: human bloodstream infection (BSI), livestock faeces (cows, pigs, poultry, and sheep), and wastewater (influent, effluent, and rivers upstream of effluent)^18,19^. All BSI isolates and a majority of non-BSI isolates were not selectively cultured for antimicrobial resistance. Species were determined with matrix-assisted laser desorption/ionization time-of-flight (MALDI-TOF). Authors used a hybrid approach (short- and long-reads) to recover reference-grade assemblies. We selected all the *E. coli* genomes where the chromosome and all plasmids were fully circularised (n= 1,044). Isolate and assembly metadata was also downloaded.

### Genome annotation

We annotated for PDSs using the tool DefenseFinder (v. 2.0.0) with default paramaters, which queries against a database of known PDSs^20–22^. DefenseFinder also denotes when a system is “complete” (“systems” output), meaning all component genes have been annotated. We only considered complete systems in this study. For ARGs, we used pre-existing annotations from AMRFinderPlus (v. 3.10.18) with default parameters^23,24^. Plasmid mobility predictions (conjugative, mobilisable, or non-mobilisable) also used pre-existing annotations from MOB-typer from MOB-suite (v. 3.03) with default parameters. We annotated for integrons using IntegronFinder 2.0 with default parameters^25^. For gene annotations we used Prodigal (v. 2.6.3)^26^ with default parameters.

### Testing for the enrichment of ARGs and PDS genes on plasmids

We first calculated the total number of plasmid genes as a fraction of the total number of genes in the genome. Similarly, we calculated the fraction of just ARGs and PDSs genes on plasmids. These proportions were compared using a Pearson’s chi-squared test with Yates’ continuity correction. We conducted this across the whole dataset (see Results), but also separately for BSI-associated and non-BSI-associated plasmids (Figure S5).

### Testing for the over-representation of PDS families on plasmids

First, for each PDS type, we calculated a binary presence/absence score for each replicon. We next wanted to generate a null distribution that accounted for some genomes containing more plasmids than others. To do this, we performed *b*=10,000 label permutations of replicon type (plasmid or chromosome) within each genome. After each permutation, we recorded the number of plasmid-labelled contigs carrying each PDS. This generated a one-sided *p*-value=(*n*+1)/*(b*+1), where *n* was the number of permutations with a plasmid count at least as extreme as observed (≥ for the plasmid-enrichment test; ≤ for the chromosome-enrichment test) and *b*=10,000. Separate *p*-values were obtained for plasmid (H_1_: observed > null) and chromosome (H_1_: observed < null) bias. To mitigate against multiple tests increasing the rate of Type I errors, we applied both the Benjamini–Hochberg (BH) and the Benjamini–Yekutieli (BY) false-discovery-rate procedures. Families with BH-adjusted q < 0.025 for the relevant one-sided test (plasmid or chromosome) were called significant; BY-adjusted q-values are also reported when q < 0.025 (providing a stricter, dependence-robust correction).

### Testing the association between ARGs and PDSs on plasmids

If two plasmids are vertical descendants, or share large portions of their sequence from a recombination event, they might contain many of the same genes. This can be problematic when testing for the association of ARGs and PDSs, as it introduces pseudo-replicates to the model. We minimised the risk of pseudo-replication by genetically dereplicating plasmids in the dataset. We first took the *k*-mer plasmid clusters generated by the original authors^17^. We then randomly selected a representative from each cluster. In total, we recovered *n*=712 genetically representative plasmids.

We next predicted the count of ARGs per plasmid with a generalised additive mixed-model (GAMM) using the mgcv library^27^. Model choice was guided by Akaike Information Criterion (AIC), which balances goodness-of-fit and parsimony. We found that a Tweedie family GAMM with a log-link family outperformed analogous GAMM/GLMMs with Poisson and negative binomial families.

The model used the fixed effects (i) plasmid PDS count, (ii) plasmid GC content (mean centred), and (iii) sampling niche (BSI-associated, livestock-associated, or wastewater-associated). We used plasmid replicon type as a random intercept to control for plasmid lineage. We also included an offset of log_10_-scaled plasmid length (bp). Originally, we also included plasmid integron count and plasmid mobility (non-mobilisable, mobilisable, conjugative) as fixed-effects, but these were dropped since they were insignificant (likelihood-ratio test *p*-value>0.05). Plasmid PDS count and GC content initially used thin-plate regression splines (*k* = 4), but by assessing the effective degrees of freedom and plots of the smooth terms, we concluded they were consistent with linearity, so the smooth terms were dropped.

Lastly, we checked for multicollinearity and correlations among predictors. For numeric-numeric fixed-effect pairs, we computed Spearman’s rank correlation, and for numeric-categorical fixed-effect pairs, we calculated the Kruskal-Wallis effect size. No issues were identified.

Of the *n*=712 genetically representative plasmids, 23% (167/712) had inconclusive plasmid replicon typing by PlasmidFinder (recorded as “NA”). Nonetheless, re-running the GAMM with these plasmids excluded did not meaningfully change the coefficient estimates (95% CIs overlapped).

### Code and data availability

The data and metadata used in this study will be stably archived on Zenodo. The code to reproduce the analysis will be made available on GitHub, and also archived on Zenodo. Supplementary File 1 contains the permutation test output. Supplementary File 2 contains the GAMM diagnostics.

## Results

### Plasmids are enriched for antimicrobial resistance genes and phage-defense system genes

We first curated a sample of *n*=1,044 *E. coli* isolates, a common human commensal and pathogen, with *n*=2,559 plasmids (see Methods). Isolates were sampled within a geographically (<60km in Oxfordshire, UK) and temporally (2008-2020) restricted frame, and represented both clinical (human bloodstream infection [BSI]) and non-clinical (livestock-associated and wastewater-associated) niches, totaling 52% (547/1,044) BSI, 42% (433/1,044) livestock-associated, and 6% (64/1,044) wastewater-associated. Livestock-associated isolates were further divided into 32.8% (142/433) cattle-associated, 28.9% (125/433) pig-associated, 11.8% (51/433) poultry-associated, and 26.5% (115/433) sheep-associated. In total, 89% (938/1,044) of isolates carried a plasmid. Isolates carried a median of 2 plasmids (range: 0-16), with the most common replicon types being small colicinogenic-type plasmids (see Methods).

We next annotated our genomes for ARGs and PDSs (see Methods). In total, we identified *n*=14,013 PDS hits with 180 distinct system types. Every chromosome carried at least one PDS, contrasted with only 41% (1,051/2,559) of plasmids. Comparatively, ARGs were less common, with 3,193 identified in total. Overall, 27% (286/1,044) of chromosomes and 20% (521/2,559) of plasmids carried at least one ARG, respectively. Of plasmids that carried at least one ARG or PDS, the three top replicon types were IncF-types: IncFIB, Col156-IncFIB-IncFII, and IncFIB-IncFIC, which together accounted for 27% (325/1,225) of that plasmid subset.

Distributions of ARGs and PDSs were variable across *E. coli* niches (Figures S1–S4). Livestock-associated plasmids (cattle, pig, poultry, and sheep) carried defense systems in 44% (475/1,080) of cases but carried ARGs in only 13% (143/1,080). Within this group, pig-associated plasmids were distinct: they encoded ARGs roughly four-fold more frequently: 25% (101/403) of pig plasmids versus 6% (42/677) of plasmids from the other three livestock groups, while their PDS prevalence remained comparable (39% [158/403] versus 47% [317/677]). Wastewater-associated plasmids showed intermediate prevalence: 35% (53/150) encoded PDSs and 19% (29/150) carried ARGs. BSI-associated plasmids exhibited the greatest combined load outside pigs, with 39% (523/1,329) containing PDSs and 26% (349/1,329) carrying ARGs.

Next, we tested whether ARGs and PDSs were enriched on plasmids versus chromosomes (Figure 1; see Methods). Of the *n*=4,999,278 annotated coding sequences in our dataset, only 3.42% (170,778/4,999,278) were found on plasmids. By contrast, 8.66% of all PDS genes (2,474/28,581) and 75.76% of all ARGs (2,419/3,193) were plasmid-encoded. Pearson’s χ^2^ tests (Yates-corrected) confirmed that both categories were highly over-represented on plasmids relative to the genomic baseline (ARGs: χ^2^ = 49,943.4, df = 1, *p* < 0.001; PDSs: χ^2^ = 2,343.8, df = 1, *p* < 0.001). The magnitude of enrichment was therefore ∼2.5-fold for PDS genes and ∼22-fold for ARGs. Surprisingly, this trend carried for both clinical and non-clinical isolates (Figure S5).

**Figure 1.**
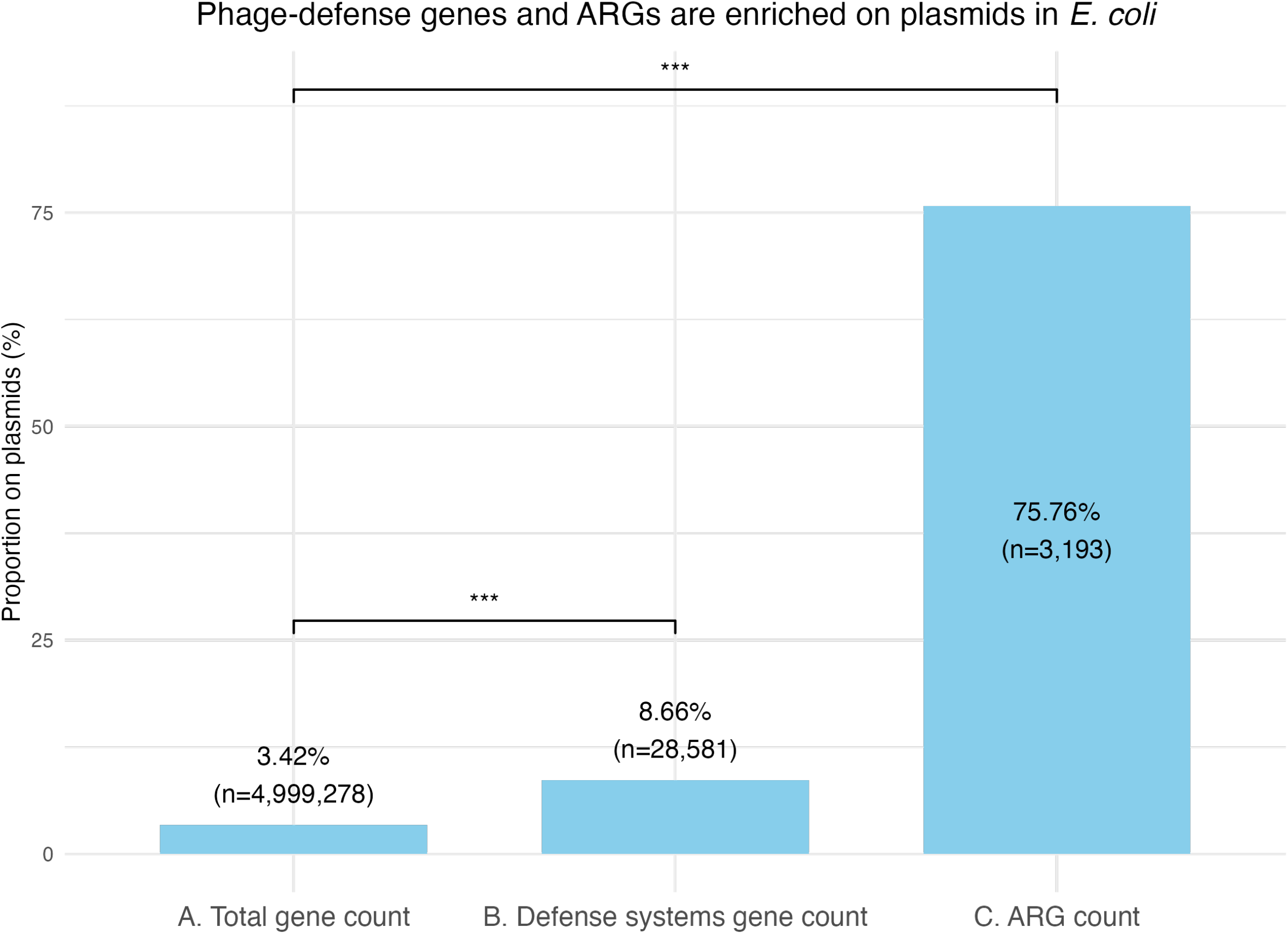
Plasmids are hotspots for ARGs and PDSs in *E. coli*. Bars show the percentage of genes in that category residing on plasmids. Both comparisons were significant at the p<0.001 level (Pearson χ^2^ tests with Yates correction).

### Plasmids and chromosomes are associated with distinct repertoires of phage-defense systems

We next explored whether the types of PDSs varied between plasmids and chromosomes. We conducted a permutation test for each PDS (see Methods) and found 8% (15/180) of the PDS families preferentially resided on plasmids (Figure 2). These 15 families made up 42.46% (1,343/3,163) of defense-system occurrences on plasmids. In contrast, 20% (36/180) PDSs were associated with chromosomes (Figure S6). Tests for the remaining 72% (129/180) PDSs were inconclusive.

**Figure 2.**
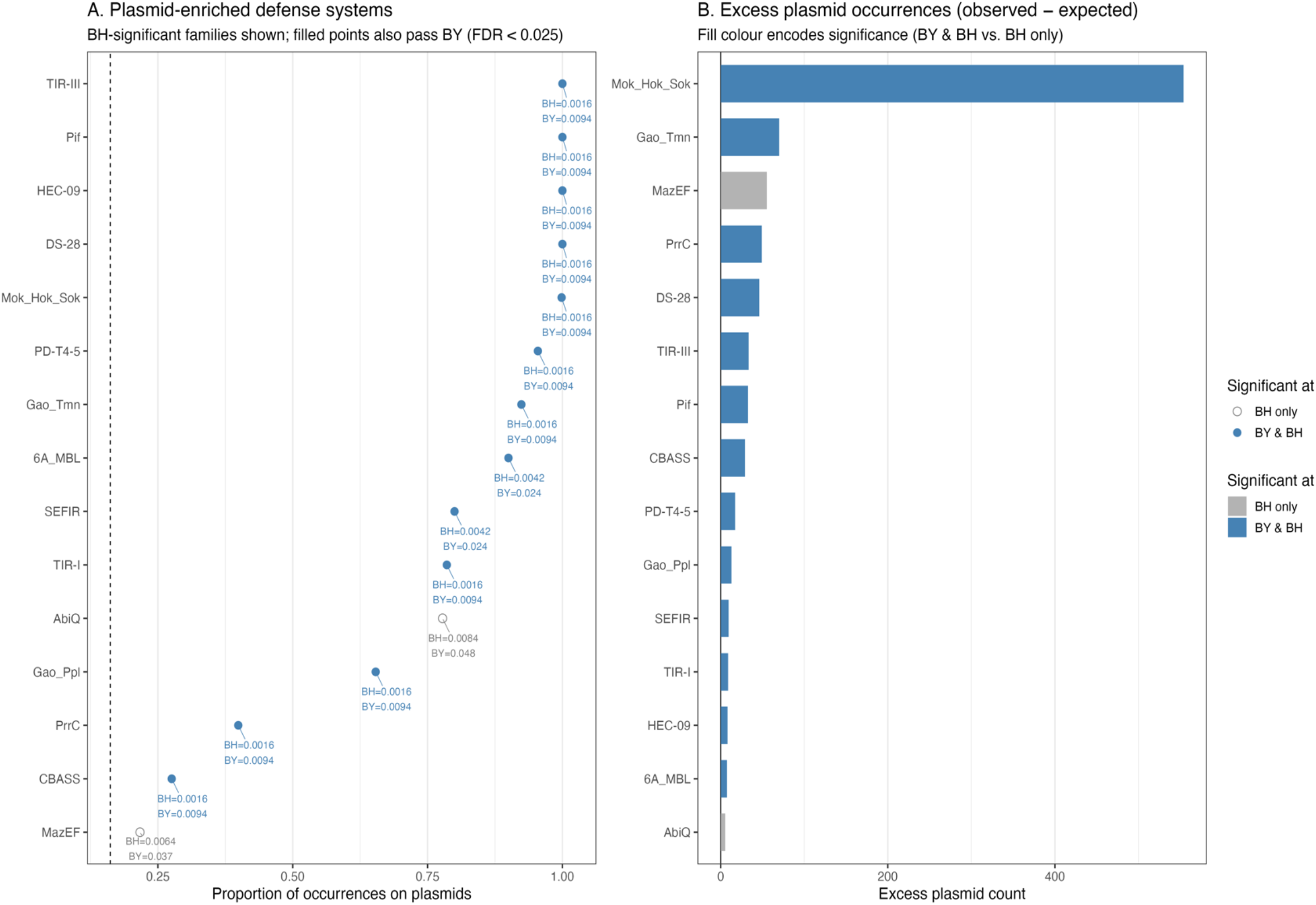
Defense systems associated with plasmids. **(A)** The x-axis shows the proportion of contigs that carried the PDS type. Each point represents a PDS type whose frequency on plasmids was greater than expected when contig labels (plasmid/chromosome) were randomly shuffled within isolates under two one-tailed permutation tests (see Methods). Open grey circles indicate p<0.05 significance under Benjamini-Hochberg (BH) correction only. Filled blue circles indicate significance under both BH and Benjamini-Yekutieli (BY) correction, which does not assume independence. The dashed line marks the genome-wide plasmid share of defence systems 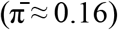. **(B)** Bars count the excess number of plasmid-labelled contigs carried by each type (“observed – expected”, where the expectation equals the type’s total contig count ×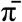). Bars are coloured the same as points in **(A)**. Only PDS families that were significantly over-represented on plasmids in the within-isolate permutation test are shown.

We found that TIR-III and Pif systems and two small uncharacterised families (HEC-09, DS-28) were exclusively associated with plasmids in this dataset. Conversely, toxin-antitoxin module MazEF and the cyclic-oligonucleotide producer CBASS were enriched more modestly (∼0.25, Figure 2A). When enrichment was quantified as absolute surplus (Figure 2B), the toxin-antitoxin complex Mok-Hok-Sok was an outlier, occurring on plasmids 661 times versus the approximately 100 occurrences expected by chance. Progressively smaller and significant surpluses were observed for the remaining systems.

### Phage-defense system count predicts ARG gene count on plasmids independent of plasmid size

Lastly, we explored the distribution of ARGs and PDSs within the genomes. We tested whether ARGs and PDSs co-occurred on the same plasmids by constructing a generalised additive mixed-model (GAMM) with a Tweedie family and log-link (see Methods). Briefly, we used plasmid ARG count as a response with fixed effects (i) plasmid PDSs count, (ii) plasmid GC content (mean centred), and (iii) sampling niche (BSI-associated, livestock-associated, or wastewater-associated). We used plasmid replicon type as a random intercept to control for plasmid lineage. To avoid pseudo-replication of plasmid sequences, we used single representatives from clusters generated by genetic similarity. We also included an offset of log_10_-scaled plasmid length (bp). Our total sample size was *n*=712 genetically representative plasmids. The model output is detailed in Table 1 and visualised in Figure 3a. Further, Figure 3b visualises the ARG-PDS co-occurrence heatmap for all plasmids (*n*=2,559) in the dataset.

**Table 1.**
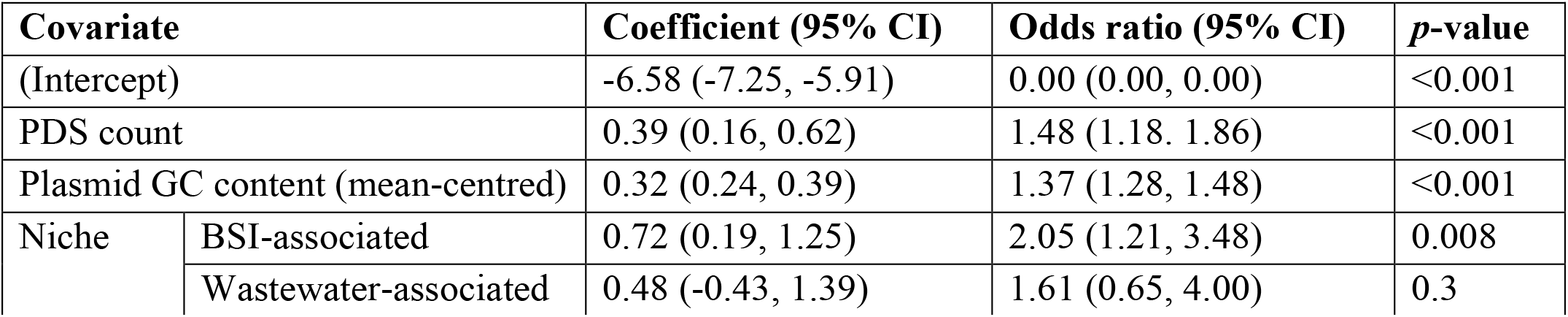
Parameter estimates for the GAMM. For niche, livestock-associated was the reference level. Odds ratios were calculated by taking the exponent.

| Covariate |  | Coefficient (95% CI) | Odds ratio (95% CI) | *p*-value |
| --- | --- | --- | --- | --- |
| (Intercept) |  | -6.58 (-7.25, -5.91) | 0.00 (0.00, 0.00) | <0.001 |
| PDS count |  | 0.39 (0.16, 0.62) | 1.48 (1.18, 1.86) | <0.001 |
| Plasmid GC content (mean-centred) |  | 0.32 (0.24, 0.39) | 1.37 (1.28, 1.48) | <0.001 |
| Niche | BSI-associated | 0.72 (0.19, 1.25) | 2.05 (1.21, 3.48) | 0.008 |
|  | Wastewater-associated | 0.48 (-0.43, 1.39) | 1.61 (0.65, 4.00) | 0.3 |

**Figure 3.**
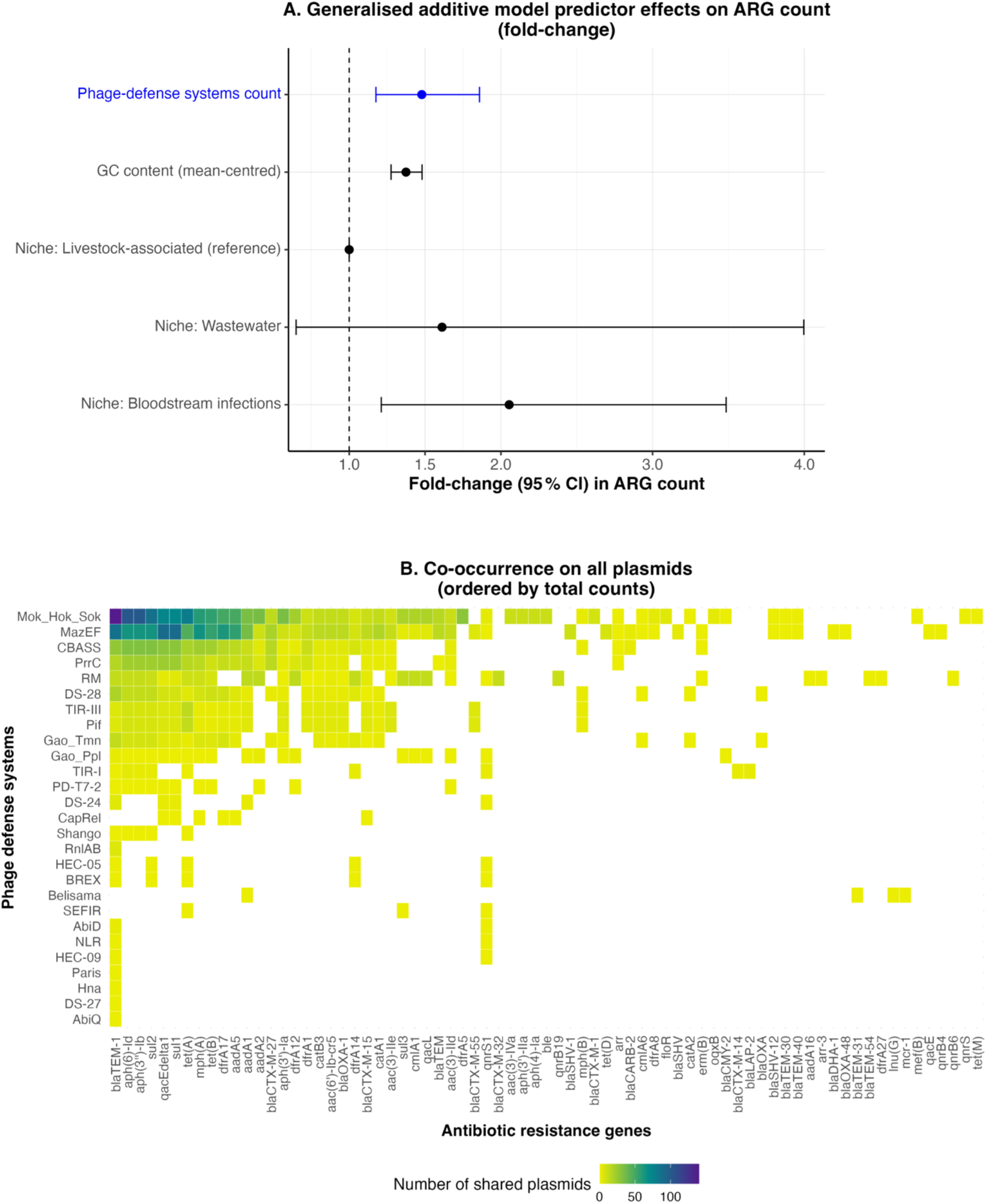
PDSs and ARGs co-occur on plasmids. **(A)** Forest plot of GAMM predictor coefficients as fold-changes with 95% confidence intervals (log_10_-scaled axis). The vertical dashed line at *x*=1 marks the reference. **(B)** Heatmap of frequency of co-localisation of ARGs and PDSs on same plasmid (*n*= 347). Cell colour indicates count.

We found that the count of PDSs was a strong positive predictor of ARG count: each additional phage-defense system increased the chance of an ARG being present on a plasmid by 48% (95% CI=[18%, 86%]; *p*<0.001). Similarly, each 1% increase in plasmid GC content above average increased ARG count by 37% (95% CI=[28%, 48%]; *p*<0.001). We found no signal from isolate sampling niche. The random-effect smooth term for plasmid replicon type was consistent with substantial non-linear variation (edf = 25.7, *F* = 1.21, *p*<0.001). Overall, the model demonstrated good fit with adjusted *R*^2^=0.40.

All isolates (1044/1044) carried a chromosomal PDS, meaning that all isolates with plasmid-borne ARG (429/1,044) had the potential for dual resistance. Additionally, of the 41% (429/1,044) of isolates carrying an ARG-positive plasmid, 88% (378/429) also carried at least one plasmid-borne PDS. In 75% (324/429) of these cases, they were both on the same plasmid (Figure 4). Further, Figure S7 shows the relative frequencies of ARGs, PDSs, and their co-localisation on plasmids, and Figure S8 shows how these frequencies vary with plasmid size in clinical versus non-clinical *E. coli*, consistent with clinical isolates, unlike non-clinical isolates, often carrying small plasmids (<13kbp) that link ARGs and PDSs.

**Figure 4.**
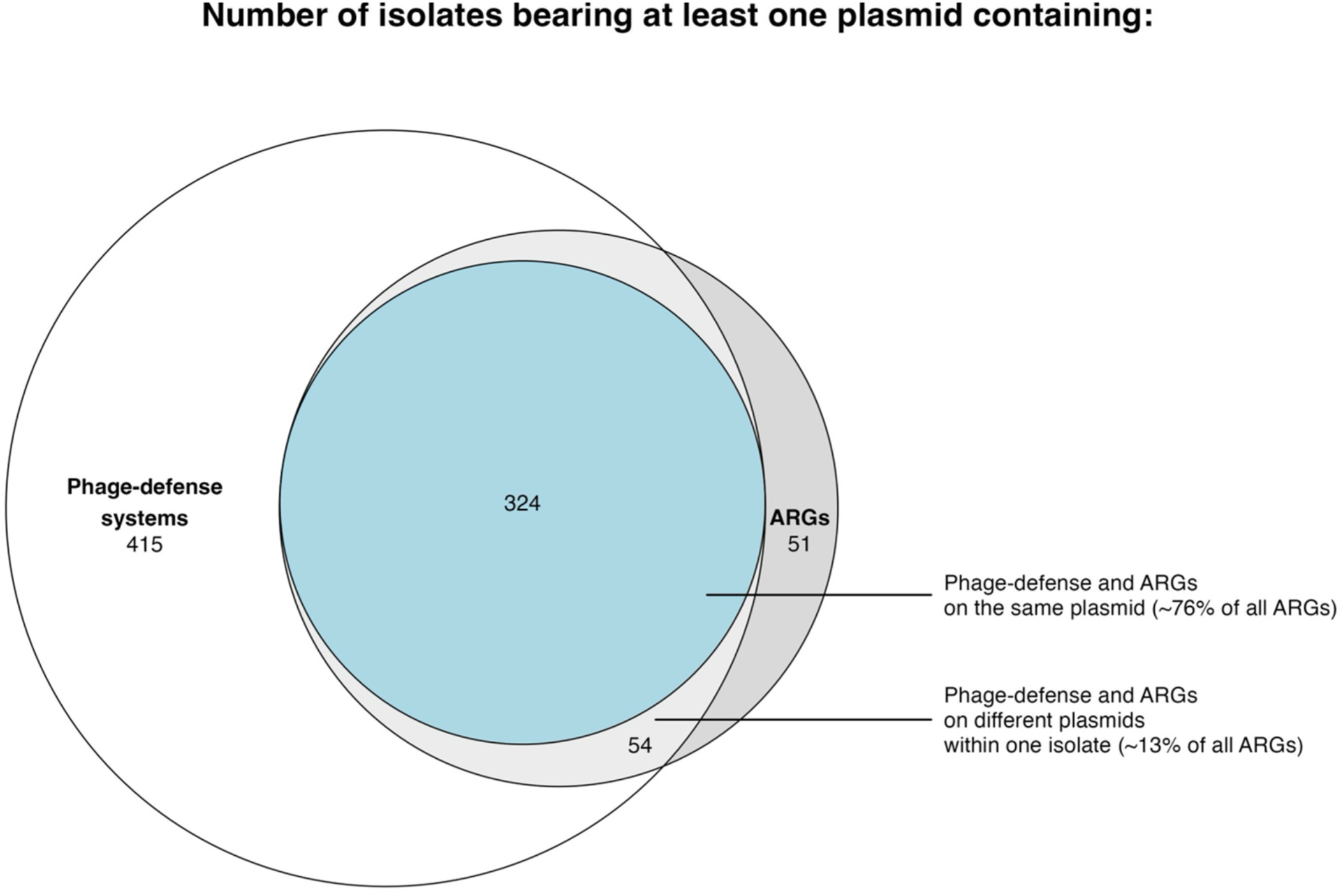
Overlap of ARGs and PDSs on plasmids within *E. coli* isolates. Euler diagram illustrating the intersection between sets of *E. coli* isolates that carried (i) at least one plasmid with a PDS, (ii) at least one plasmid with an ARG, and (iii) at least one plasmid with both an ARG and a PDS. The size of each ellipse is proportional to the set size. The total sample size was *n*=884 isolates.

## Discussion

We have analysed the distribution of ARGs and PDSs in a diverse sample of *E. coli* genomes. We demonstrate that plasmids are enriched for both types of features, disproportionately carry specific PDS families compared to the chromosome, and physically link ARGs and PDSs in the genome. Overall, our findings suggest a fundamental role for plasmids in the dissemination of PDSs on top of known associations with ARGs.

Our study has limitations. Whilst *E. coli* represents a predominant clinical pathogen, plasmids are widespread across bacterial taxa and frequently cross species boundaries^28^. Future work should expand these modelling approaches to incorporate a broader range of bacterial hosts. We also did not model the co-occurrence of specific ARGs and PDSs. Capturing such associations requires modelling frameworks that account for both vertical inheritance and horizontal transfer events (such as those mediated by transposons) as well as methods capable of resolving multivariate networks of association. Without such controls, pairwise tests risk conflating indirect correlations with direct genetic linkage, as well as inflating Type I errors^29^.

The co-localisation of ARGs and PDSs on plasmids can facilitate joint horizontal transfer and coordinated regulation. We hypothesise that plasmids are an ideal platform for co-localising ARGs and PDSs due to their variable copy number and potential for rapid evolution. In clinical isolates, multicopy plasmids carrying *bla*_TEM-1_ can amplify phenotypic resistance and accelerate adaptive evolution of the gene^30,31^. In our dataset, *bla*_TEM-1_ was often co-localised with toxin-antitoxin systems such as RnlAB, PARIS, and AbiQ^32–34^on small (>13kb) colicinogenic-type plasmids in the BSI-associated isolates. Analogously, plasmids carrying PDSs may experience rapid and fluctuating selection in response to episodic lytic phage attacks, favouring those that can quickly up or downregulate their PDSs^35,36^.

Phage pressure alone can maintain mobile genetic elements carrying ARGs even in the absence of antibiotics. For example, in *Vibrio cholerae*, episodic phage challenge increased conjugation frequencies of integrative conjugative elements carrying both ARGs and PDSs^14^. In another study, *E. coli* IncF plasmids carrying antibiotic resistance reduced susceptibility to coliphage infection and persisted for around 10 days without antibiotic selection^37^. This suggests that simply curbing antibiotic use may be insufficient to reverse the selection for antibiotic and/or phage resistance. Future work should monitor plasmid gene frequencies, copy-number dynamics, and host-fitness effects under isolated antibiotic, isolated phage, and combined selection regimes, ideally incorporating varied dosing schedules and temporal patterns.

## Funding

This work was supported by the UKRI Frontiers grant (EP/Y031067/1) to RCM. WS is supported by the Wellcome Trust grant (218514/Z/19/Z), Merck Sharp and Dohme Corp., and Janssen Pharmaceutica NV. PJ is supported by funding from the Biotechnology and Biological Sciences Research Council UKRI-BBSRC grant (BB/T008784/1).

## Acknowledgements

The authors thank Rachel Wheatley and Liam P. Shaw for the valuable help and discussions.

## Conflicting interests

The authors declare no conflicting interests.

## Notes

### Competing Interest Statement

The authors have declared no competing interest.

### Summary of Updates

We have revised the abstract to describe the results more accurately.

## Bibliography

1. Koonin, E. V., Makarova, K. S. & Wolf, Y. I. Evolutionary Genomics of Defense Systems in Archaea and Bacteria. Annu Rev Microbiol 71, (2017).

2. Coluzzi, C., Garcillán-Barcia, M. P., De La Cruz, F. & Rocha, E. P. C. Evolution of Plasmid Mobility: Origin and Fate of Conjugative and Nonconjugative Plasmids. Mol Biol Evol 39, (2022).

3. Egido, J. E., Costa, A. R., Aparicio-Maldonado, C.Haas, P.-J. & Brouns, S. J. J. Mechanisms and clinical importance of bacteriophage resistance. FEMS Microbiol Rev 46, (2022).

4. Partridge, S. R., Kwong, S. M., Firth, N. & Jensen, S. O. Mobile genetic elements associated with antimicrobial resistance. Clinical Microbiology Reviews vol. 31 Preprint at 10.1128/CMR.00088-17 (2018).

5. Vogwill, T. & Maclean, R. C. The genetic basis of the fitness costs of antimicrobial resistance: A meta-analysis approach. Evol Appl 8, (2015).

6. Siedentop, B., Rüegg, D., Bonhoeffer, S. & Chabas, H. My host’s enemy is my enemy: plasmids carrying CRISPR-Cas as a defence against phages. Proceedings of the Royal Society B: Biological Sciences 291, (2024).

7. Picton, D. M. et al. The phage defence island of a multidrug resistant plasmid uses both BREX and type IV restriction for complementary protection from viruses. Nucleic Acids Res 49, (2021).

8. Vassallo, C. N., Doering, C. R., Littlehale, M. L., Teodoro, G. I. C. & Laub, M. T. A functional selection reveals previously undetected anti-phage defence systems in the E. coli pangenome. Nat Microbiol 7, (2022).

9. Leroux, M. & Laub, M. T. Toxin-Antitoxin Systems as Phage Defense Elements. Annual Review of Microbiology vol. 76 Preprint at 10.1146/annurev-micro-020722-013730 (2022).

10. Wales, A. D. & Davies, R. H. Co-selection of resistance to antibiotics, biocides and heavy metals, and its relevance to foodborne pathogens. Antibiotics vol. 4 Preprint at 10.3390/antibiotics4040567 (2015).

11. Pal, C., Bengtsson-Palme, J., Kristiansson, E. & Larsson, D. G. J. Co-occurrence of resistance genes to antibiotics, biocides and metals reveals novel insights into their co-selection potential. BMC Genomics 16, (2015).

12. Mahata, T. et al. Gamma-Mobile-Trio systems define a new class of mobile elements rich in bacterial defensive and offensive tools. bioRxiv (2024).

13. Botelho, J. Defense systems are pervasive across chromosomally integrated mobile genetic elements and are inversely correlated to virulence and antimicrobial resistance. Nucleic Acids Res 51, (2023).

14. LeGault, K. N. et al. Temporal shifts in antibiotic resistance elements govern phagepathogen conflicts. Science (1979) 373, (2021).

15. BiomX, Inc. BiomX Announces Positive Topline Results from Phase 2 Trial Evaluating BX211 for the Treatment of Diabetic Foot Osteomyelitis (DFO). https://ir.biomx.com/news-events/press-releases/detail/130/biomx-announces-positivetopline-results-from-phase-2-trial.

16. Chan, B. K. et al. Personalized inhaled bacteriophage therapy for treatment of multidrug-resistant Pseudomonas aeruginosa in cystic fibrosis. Nat Med 31, 1494–1501 (2025).

17. Matlock, W. et al. Enterobacterales plasmid sharing amongst human bloodstream infections, livestock, wastewater, and waterway niches in Oxfordshire, UK. Elife 12, (2023).

18. Shaw, L. P. et al. Niche and local geography shape the pangenome of wastewater-and livestock-associated Enterobacteriaceae. Sci Adv 7, (2021).

19. Lipworth, S. et al. Ten-year longitudinal molecular epidemiology study of Escherichia coli and Klebsiella species bloodstream infections in Oxfordshire, UK. Genome Med 13, (2021).

20. Tesson, F. et al. Systematic and quantitative view of the antiviral arsenal of prokaryotes. Nat Commun 13, (2022).

21. Néron, B. et al. MacSyFinder v2: Improved modelling and search engine to identify molecular systems in genomes. Peer Community Journal 3, (2023).

22. Tesson, F. et al. A Comprehensive Resource for Exploring Antiphage Defense: DefenseFinder Webservice,Wiki and Databases. Peer Community Journal 4, (2024).

23. Feldgarden, M. et al. AMRFinderPlus and the Reference Gene Catalog facilitate examination of the genomic links among antimicrobial resistance, stress response, and virulence. Sci Rep 11, (2021).

24. Robertson, J. & Nash, J. H. E. MOB-suite: software tools for clustering, reconstruction and typing of plasmids from draft assemblies. Microb Genom 4, (2018).

25. Néron, B. et al. IntegronFinder 2.0: Identification and Analysis of Integrons across Bacteria, with a Focus on Antibiotic Resistance in Klebsiella. Microorganisms 10, (2022).

26. Hyatt, D. et al. Prodigal: Prokaryotic gene recognition and translation initiation site identification. BMC Bioinformatics 11, (2010).

27. Pedersen, E. J., Miller, D. L., Simpson, G. L. & Ross, N. Hierarchical generalized additive models in ecology: An introduction with mgcv. PeerJ 2019, (2019).

28. Redondo-Salvo, S. et al. Pathways for horizontal gene transfer in bacteria revealed by a global map of their plasmids. Nat Commun 11, (2020).

29. Kim, P. J. & Price, N. D. Genetic co-occurrence network across sequenced microbes. PLoS Comput Biol 7, (2011).

30. San Millan, A., Escudero, J. A., Gifford, D. R., Mazel, D. & MacLean, R. C. Multicopy plasmids potentiate the evolution of antibiotic resistance in bacteria. Nat Ecol Evol 1, 0010 (2016).

31. Matlock, W. et al. E. coli phylogeny drives co-amoxiclav resistance through variable expression of blaTEM-1. Preprint at 10.1101/2024.08.12.607562 (2024).

32. Deep, A., Liang, Q., Enustun, E., Pogliano, J. & Corbett, K. D. Architecture and infection-sensing mechanism of the bacterial PARIS defense system. (2024) doi:10.1101/2024.01.02.573835.

33. Koga, M., Otsuka, Y., Lemire, S. & Yonesaki, T. Escherichia coli rnlA and rnlB compose a novel toxin-antitoxin system. Genetics 187, (2011).

34. Pecota, D. C. & Wood, T. K. Exclusion of T4 phage by the hok/sok killer locus from plasmid R1. J Bacteriol 178, (1996).

35. Hall, A. R., Scanlan, P. D., Morgan, A. D. & Buckling, A. Host-parasite coevolutionary arms races give way to fluctuating selection. Ecol Lett 14, 635–642 (2011).

36. Betts, A., Gifford, D. R., MacLean, R. C. & King, K. C. Parasite diversity drives rapid host dynamics and evolution of resistance in a bacteria-phage system. Evolution (N Y) 70, 969–978 (2016).

37. Montelongo Hernandez, C., Putonti, C. & Wolfe, A. J. Urinary Plasmids Reduce Permissivity to Coliphage Infection. Microbiol Spectr 11, (2023).

