## Supplementary Figures for "Plasmids link antibiotic resistance genes and phage defense systems in *E. coli*"

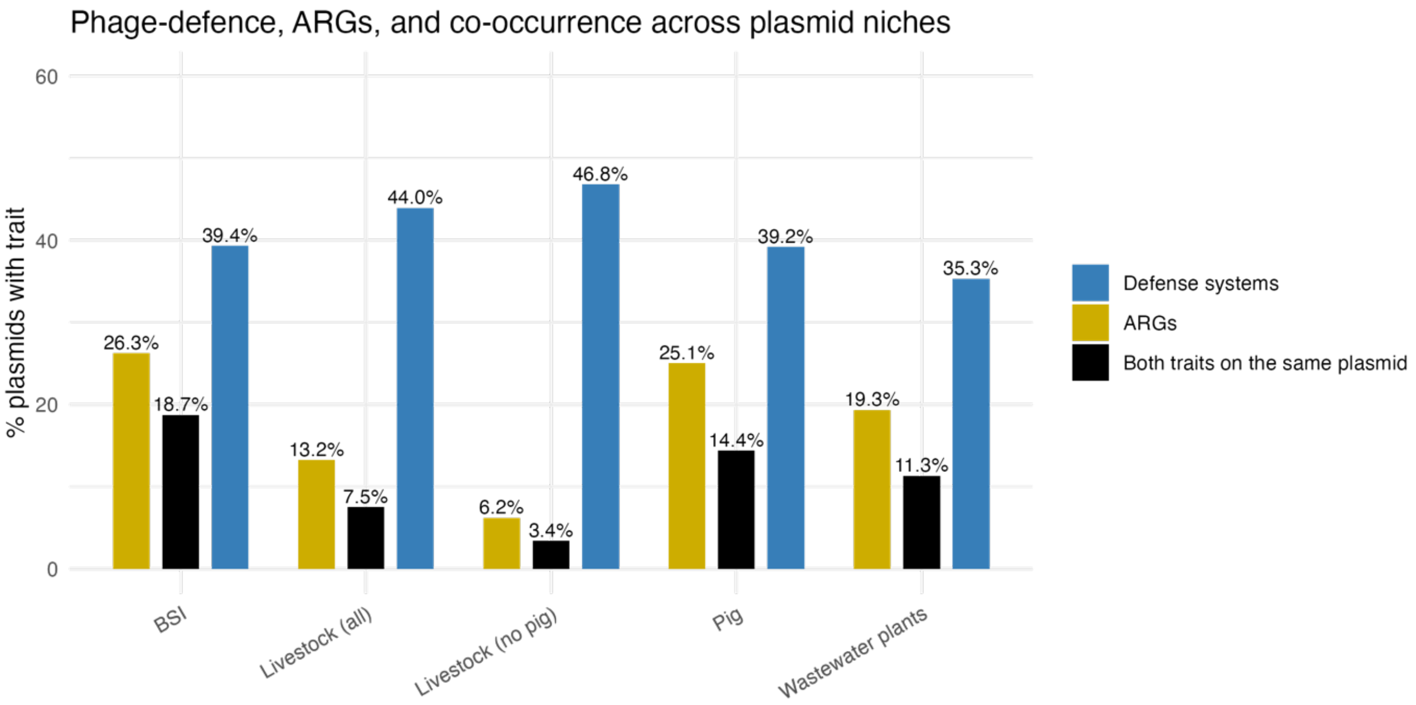


**Supplementary Figure S1. Prevalence of phage-defence systems, antibiotic-resistance genes (ARGs) and their co-occurrence in *E. coli* plasmids (n=2,559) across five ecological/clinical niches.** For each niche, bars indicate the percentage of plasmids that carry at least one phage-defence system (blue), at least one ARG (gold) or both traits on the same plasmid (black). Defence systems are consistently abundant (~35-47 %) in every environment, whereas ARGs show marked heterogeneity – highest in bloodstream-infection (BSI, 26.3%) and pig plasmids (25.1%), intermediate in wastewater-plant isolates (19.3%), and minimal in livestock (no pig) plasmids (6.2%). Plasmids harbouring both traits concentrate in BSI (18.7%) and pig (14.4%) niches, dropping to <5 % in livestock without pigs. The data highlight phage predation as a pervasive selective force across habitats, while antibiotic pressure – and thus ARG acquisition – is niche-specific and most pronounced in clinical and pig farm settings.

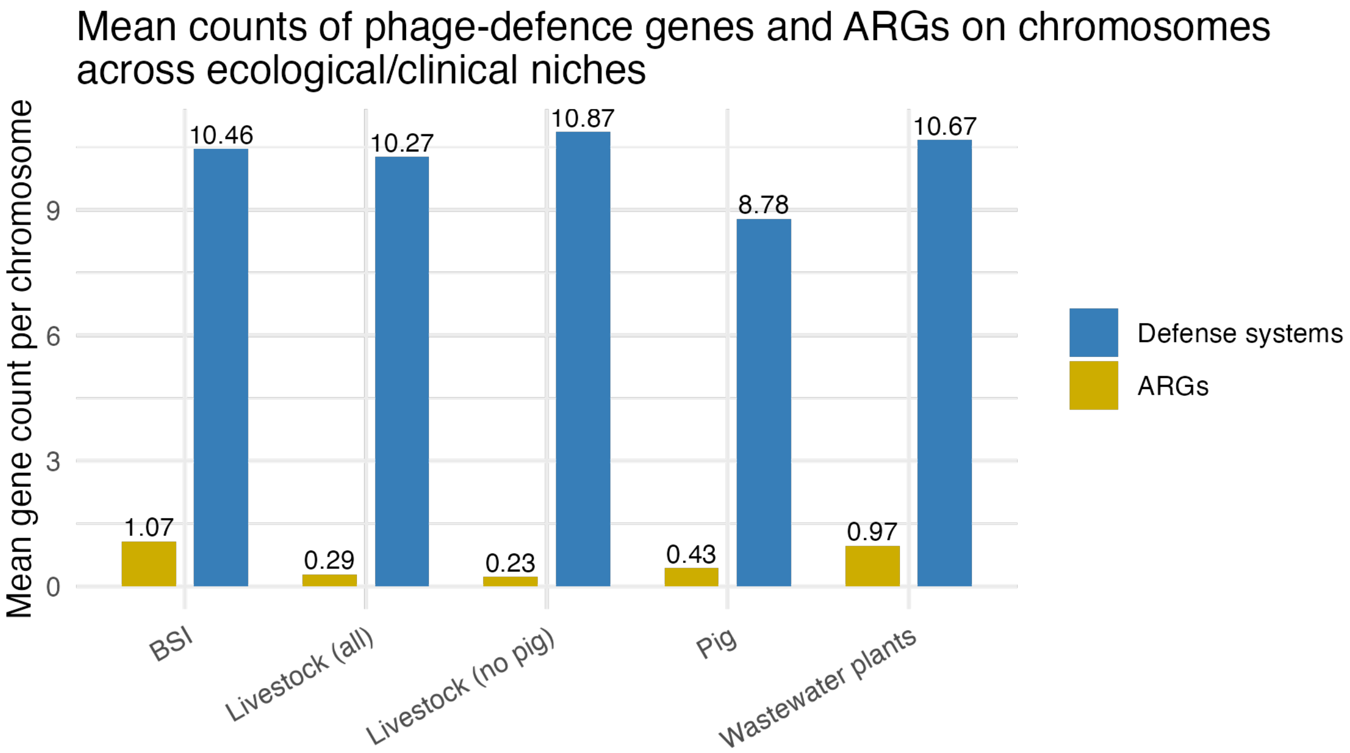


**Supplementary Figure S2. Mean copy number of phage-defence loci versus antibiotic-resistance genes (ARGs) on *E. coli* chromosomes (n=1,044) stratified by five ecological/clinical niches.** Blue bars indicate the average number of phage-defence systems per chromosome; gold bars show the corresponding mean for ARGs. Across all environments, chromosomes carry a large and uniform defence repertoire (8-11 loci). In contrast, chromosomal ARG content is consistently low (< 1 gene on average), peaking in bloodstream-infection (BSI) and wastewater-plant isolates (0.6-0.7) and falling to ≤ 0.3 in livestock-associated chromosomes. The ~15-fold excess of defence over ARG loci highlights phage predation as a pervasive selective force shaping the core genome, while chromosomal incorporation of antibiotic-resistance determinants remains rare and niche-specific.

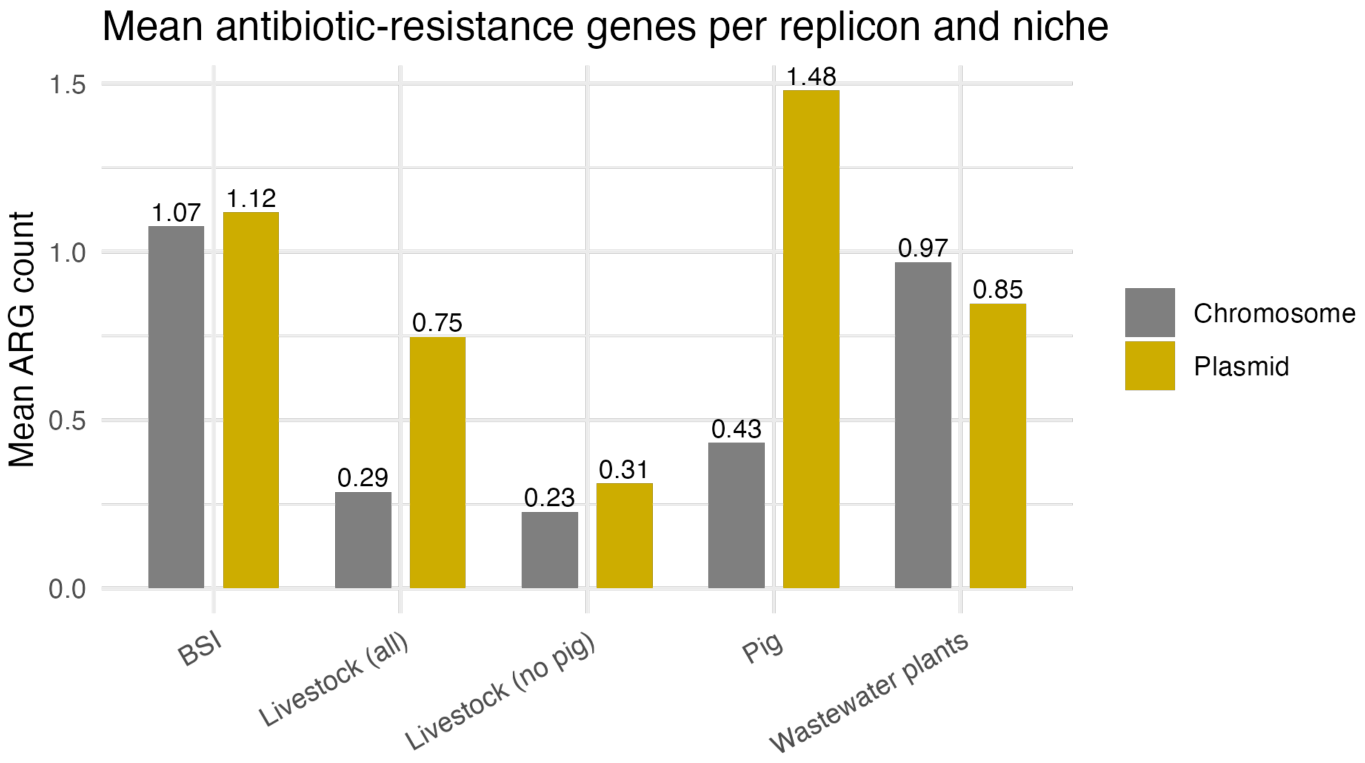


**Supplementary Figure S3. Mean burden of antibiotic-resistance genes (ARGs) on plasmids versus chromosomes in five *E. coli* niches (plasmids n=2,559; chromosomes n=1,044).** Gold bars show the average number of ARGs per plasmid; grey bars the average per chromosome. Plasmids consistently exceed chromosomes in ARG content, with the largest differential in pig isolates (1.48 vs 0.43 genes) and wastewater-plant samples (0.85 vs 0.97 genes). Livestock chromosomes are almost ARG-free (≤ 0.30 genes on average), while plasmids from the same hosts retain a modest resistance load (0.75 genes). Bloodstream infection (BSI) genomes carry the heaviest chromosomal ARG burden (1.07), yet plasmids still marginally outpace them (1.12). These patterns underscore plasmids as the primary vector for resistance dissemination, particularly in clinical and intensive-pig environments, whereas chromosomal acquisition of ARGs remains limited in most non-clinical settings.

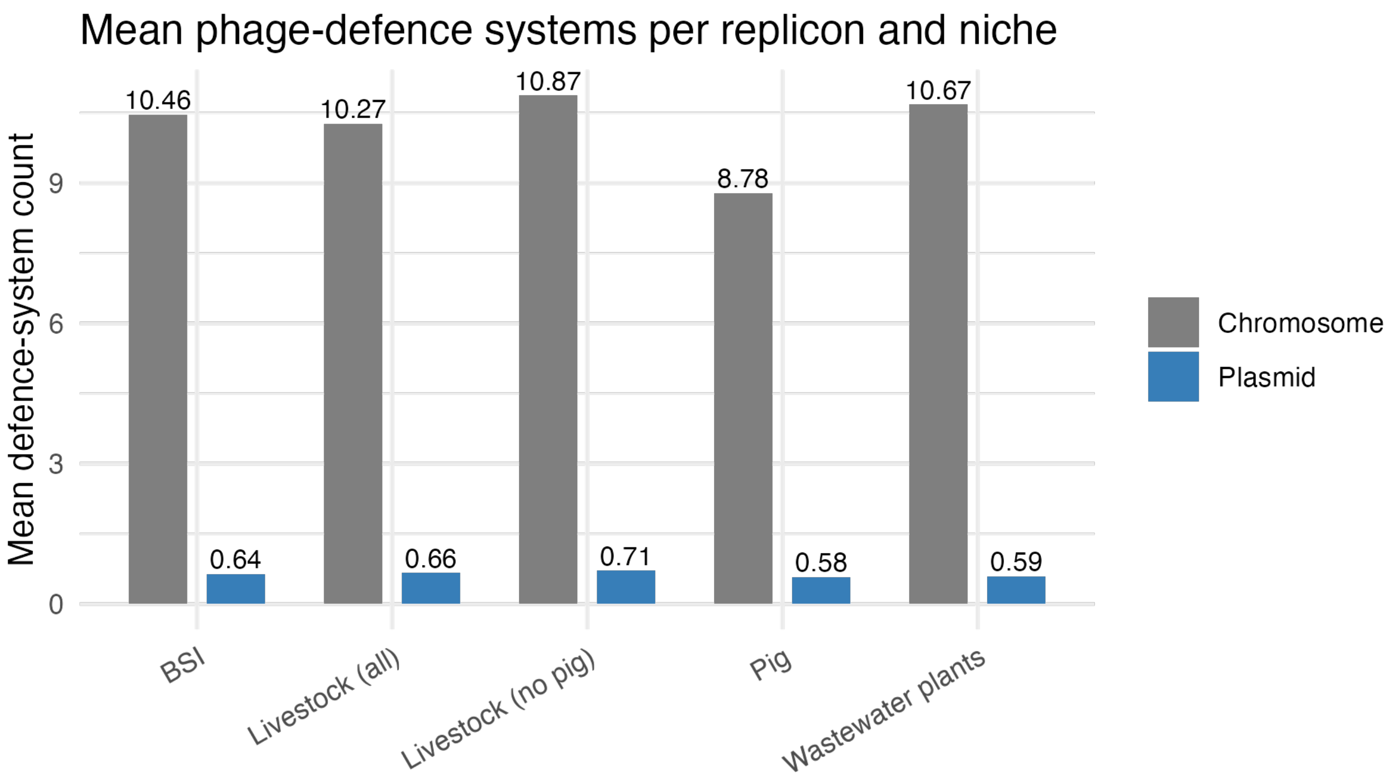


**Supplementary Figure S4. Mean copy number of phage-defence systems on plasmids versus chromosomes across five *E. coli* niches (plasmids n=2,559; chromosomes n=1,044).** Grey bars indicate the average number of defence loci per chromosome; blue bars show the corresponding mean per plasmid. Chromosomes carry a large and stable repertoire (~9-11 systems) in every niche, whereas plasmids harbour ≤ 0.7 systems on average, with only minor niche-to-niche variation.

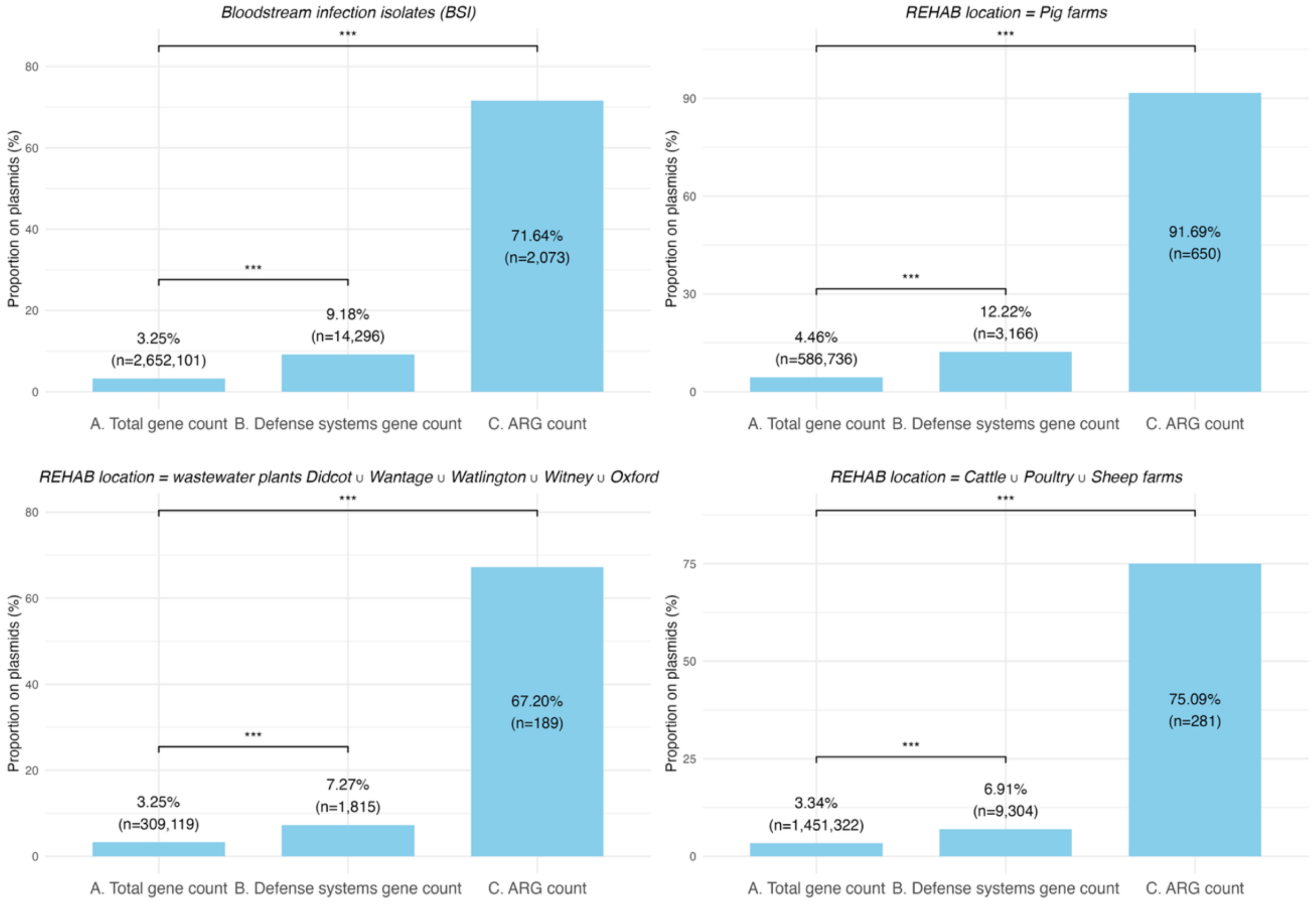


**Supplementary Figure S5. Fraction of E. coli genes carried on plasmids in four epidemiological settings.** Stacked panels compare the proportion of genes located on plasmids for three functional categories – (A) total annotated genes, (B) phage-defence system genes, and (C) antimicrobial-resistance genes (ARGs) – in isolates from (top-left) human bloodstream infections (BSI), (top-right) pig farms, (bottom-left) municipal wastewater treatment plants (Didcot, Wantage, Watlington, Witney and Oxford) and (bottom-right) cattle, poultry and sheep farms. Bar heights give the percentage of category-specific genes found on plasmids; absolute gene counts are shown in parentheses inside each bar. Horizontal brackets indicate pairwise $\chi^2$ tests: *** $P < 0.001$ for ARGs vs. total genes and for defence-system genes vs. total genes in every panel, and *** $P < 0.001$ for ARGs vs. defence-system genes. Across all environments, ARGs are overwhelmingly plasmid-borne (63-92 %), defence-system genes are enriched but to a lesser extent (6-12 %), and only ~3-4 % of the average gene complement is plasmid encoded, underscoring the central role of plasmids in mobilising both antimicrobial resistance and phage-defence functions.

(A)

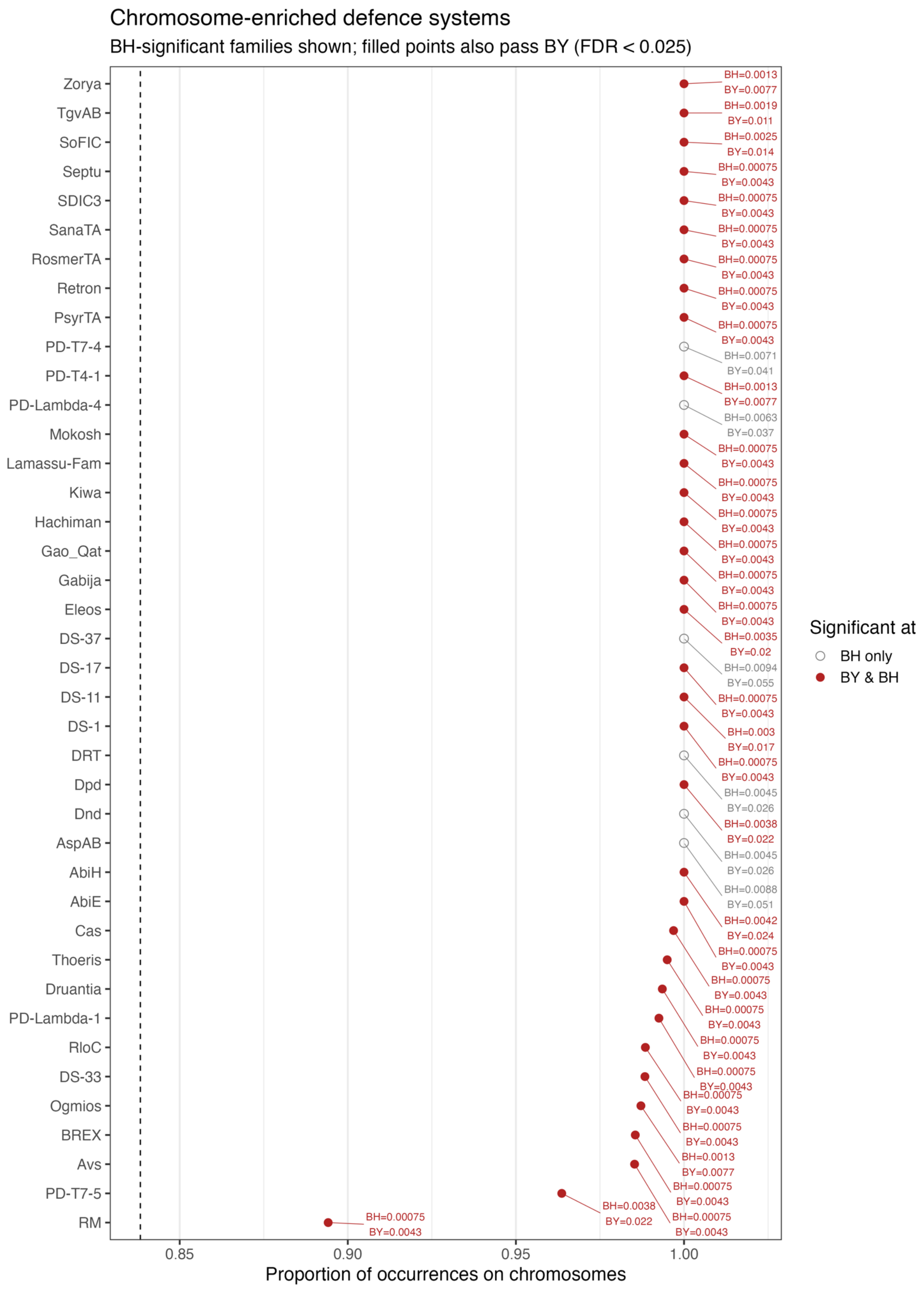

Chromosome-enriched defence systems
BH-significant families shown; filled points also pass BY (FDR < 0.025)
Zorya
TgvAB
SoFIC
Septu
SDIC3
SanaTA
RosmerTA
Retron
PsyrTA
PD-T7-4
PD-T4-1
PD-Lambda-4
Mokosh
Lamassu-Fam
Kiwa
Hachiman
Gao_Qat
Gabija
Eleos
DS-37
DS-17
DS-11
DS-1
DRT
Dpd
Dnd
AspAB
AbiH
AbiE
Cas
Thoeris
Druantia
PD-Lambda-1
RloC
DS-33
Ogmios
BREX
Avs
PD-T7-5
RM
BH=0.0013 BY=0.0077
BH=0.0019 BY=0.011
BH=0.0025 BY=0.014
BH=0.00075 BY=0.0043
BH=0.00075 BY=0.0043
BH=0.00075 BY=0.0043
BH=0.00075 BY=0.0043
BH=0.00075 BY=0.0043
BH=0.00075 BY=0.0043
BH=0.0071 BY=0.041
BH=0.0013 BY=0.0077
BH=0.0063 BY=0.037
BH=0.00075 BY=0.0043
BH=0.00075 BY=0.0043
BH=0.00075 BY=0.0043
BH=0.00075 BY=0.0043
BH=0.00075 BY=0.0043
BH=0.00075 BY=0.0043
BH=0.0035 BY=0.02
BH=0.0094 BY=0.055
BH=0.00075 BY=0.0043
BH=0.003 BY=0.017
BH=0.00075 BY=0.0043
BH=0.0045 BY=0.026
BH=0.0038 BY=0.022
BH=0.0045 BY=0.026
BH=0.0088 BY=0.051
BH=0.0042 BY=0.024
BH=0.00075 BY=0.0043
BH=0.00075 BY=0.0043
BH=0.00075 BY=0.0043
BH=0.00075 BY=0.0043
BH=0.00075 BY=0.0043
BH=0.00075 BY=0.0043
BH=0.00075 BY=0.0043
BH=0.0013 BY=0.0077
BH=0.00075 BY=0.0043
BH=0.0038 BY=0.022
BH=0.00075 BY=0.0043
BH=0.00075 BY=0.0043
0.85
0.90
0.95
1.00
Proportion of occurrences on chromosomes
Significant at
BH only
BY & BH

(B)

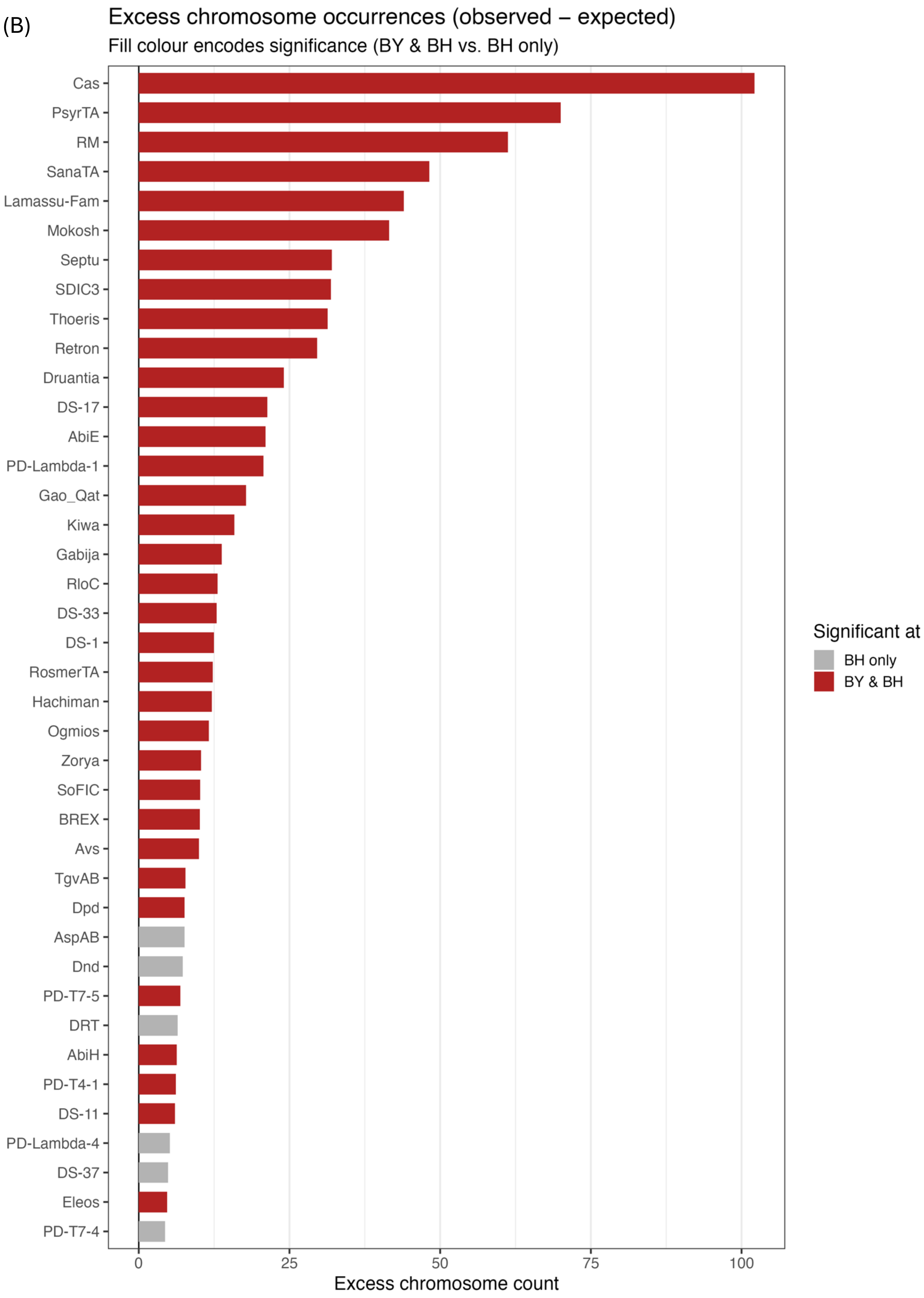

Excess chromosome occurrences (observed – expected)
Fill colour encodes significance (BY & BH vs. BH only)
Cas
PsyrTA
RM
SanaTA
Lamassu-Fam
Mokosh
Septu
SDIC3
Thoeris
Retron
Druantia
DS-17
AbiE
PD-Lambda-1
Gao_Qat
Kiwa
Gabija
RloC
DS-33
DS-1
RosmerTA
Hachiman
Ogmios
Zorya
SoFIC
BREX
Avs
TgvAB
Dpd
AspAB
Dnd
PD-T7-5
DRT
AbiH
PD-T4-1
DS-11
PD-Lambda-4
DS-37
Eleos
PD-T7-4
0
25
50
75
100
Excess chromosome count
Significant at
BH only
BY & BH

**Supplementary Figure S6. (A) Defence-system families significantly enriched on chromosomes.** Each point denotes a family whose observed frequency on chromosomal contigs exceeds the null expectation when contig labels are permuted within each isolate (10 000 permutations; Benjamini–Hochberg FDR < 0.025). Open grey circles mark families significant only by BH, whereas filled red circles also satisfy the stricter Benjamini–Yekutieli (BY) correction (legend, right). Numeric labels beneath each point now show both BH and BY q-values (first and second line, respectively). The x-axis gives the proportion of contigs carrying the family that are chromosomal; the dashed vertical line indicates the genome-wide chromosomal baseline ($1 - \bar{\pi} \approx 0.84$). Families are ordered by increasing chromosomal share. While nearly all significant families are almost exclusively chromosomal (e.g. Cas, Septu, Zorya, TgvAB with > 99 % chromosomal occurrence), the restriction–modification (RM) system and PD-T7-5 show more moderate enrichment. **(B) Absolute surplus of chromosomal occurrences for permutation-significant defence-system families.** Bars show observed - expected counts (expected = total occurrences × chromosomal baseline). Bars are coloured as in panel A: red for families significant by both BH and BY, grey for BH-only, and numerical q-values are omitted for clarity. The classical Cas adaptive-immunity operon exhibits the largest surplus (~105 excess chromosomal hits), followed by pan-immune PsyrTA, RM and several toxin-antitoxin or DNA-cleavage systems (e.g. SanaTA, Lamassu-Fam, Mokosh, Septu, SDIC3), each showing 30-60 excess occurrences. A gradient of smaller yet significant surpluses extends through more niche systems, underscoring that chromosomal residency varies continuously among *E. coli* defence elements.

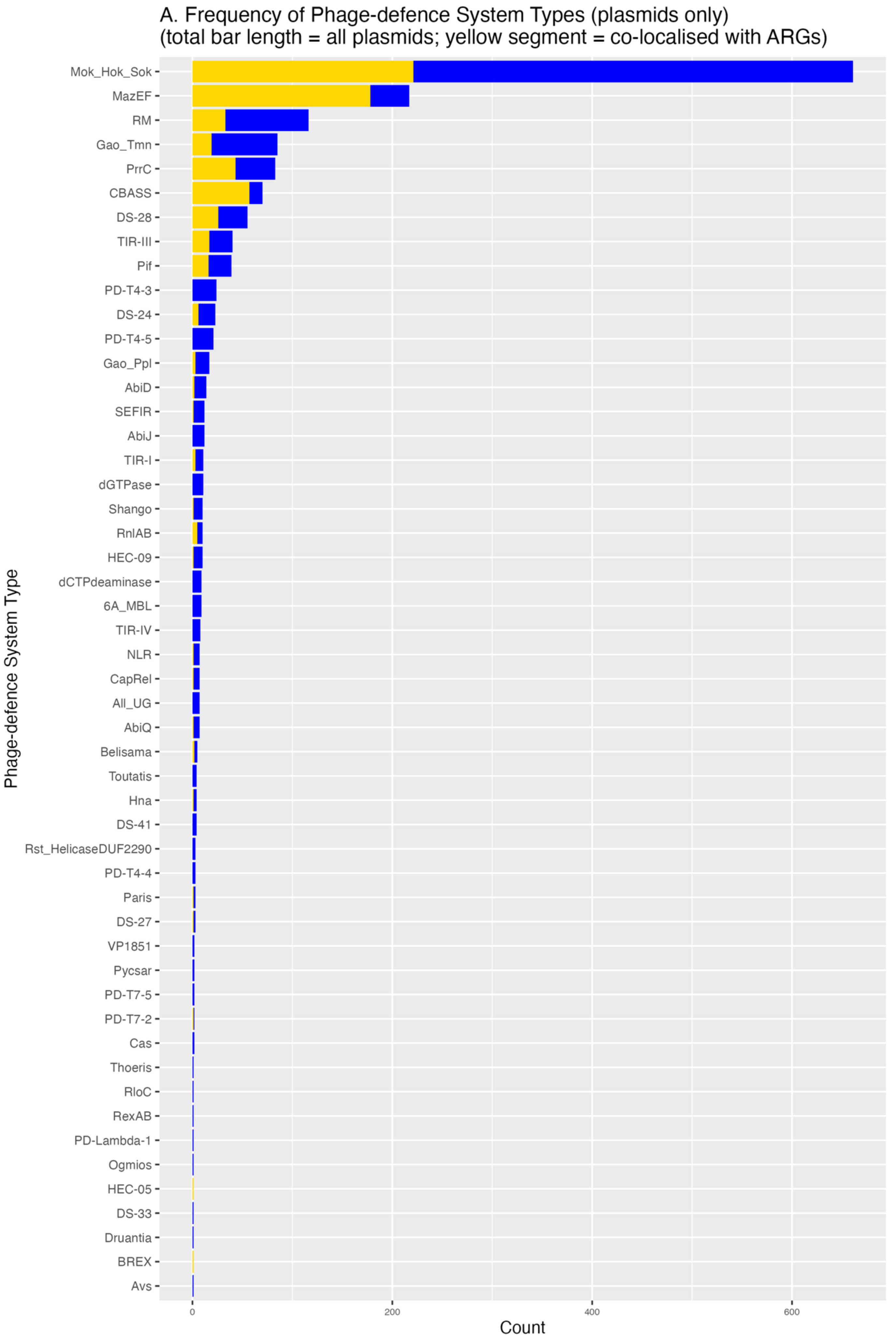
A. Frequency of Phage-defence System Types (plasmids only)
(total bar length = all plasmids; yellow segment = co-localised with ARGs)
Phage-defence System Type
Count
0
200
400
600
Mok_Hok_Sok
MazEF
RM
Gao_Tmn
PrrC
CBASS
DS-28
TIR-III
Pif
PD-T4-3
DS-24
PD-T4-5
Gao_Ppl
AbiD
SEFIR
AbiJ
TIR-I
dGTPase
Shango
RnlAB
HEC-09
dCTPdeaminase
6A_MBL
TIR-IV
NLR
CapRel
All_UG
AbiQ
Belisama
Toutatis
Hna
DS-41
Rst_HelicaseDUF2290
PD-T4-4
Paris
DS-27
VP1851
Pycsar
PD-T7-5
PD-T7-2
Cas
Thoeris
RloC
RexAB
PD-Lambda-1
Ogmios
HEC-05
DS-33
Druantia
BREX
Avs

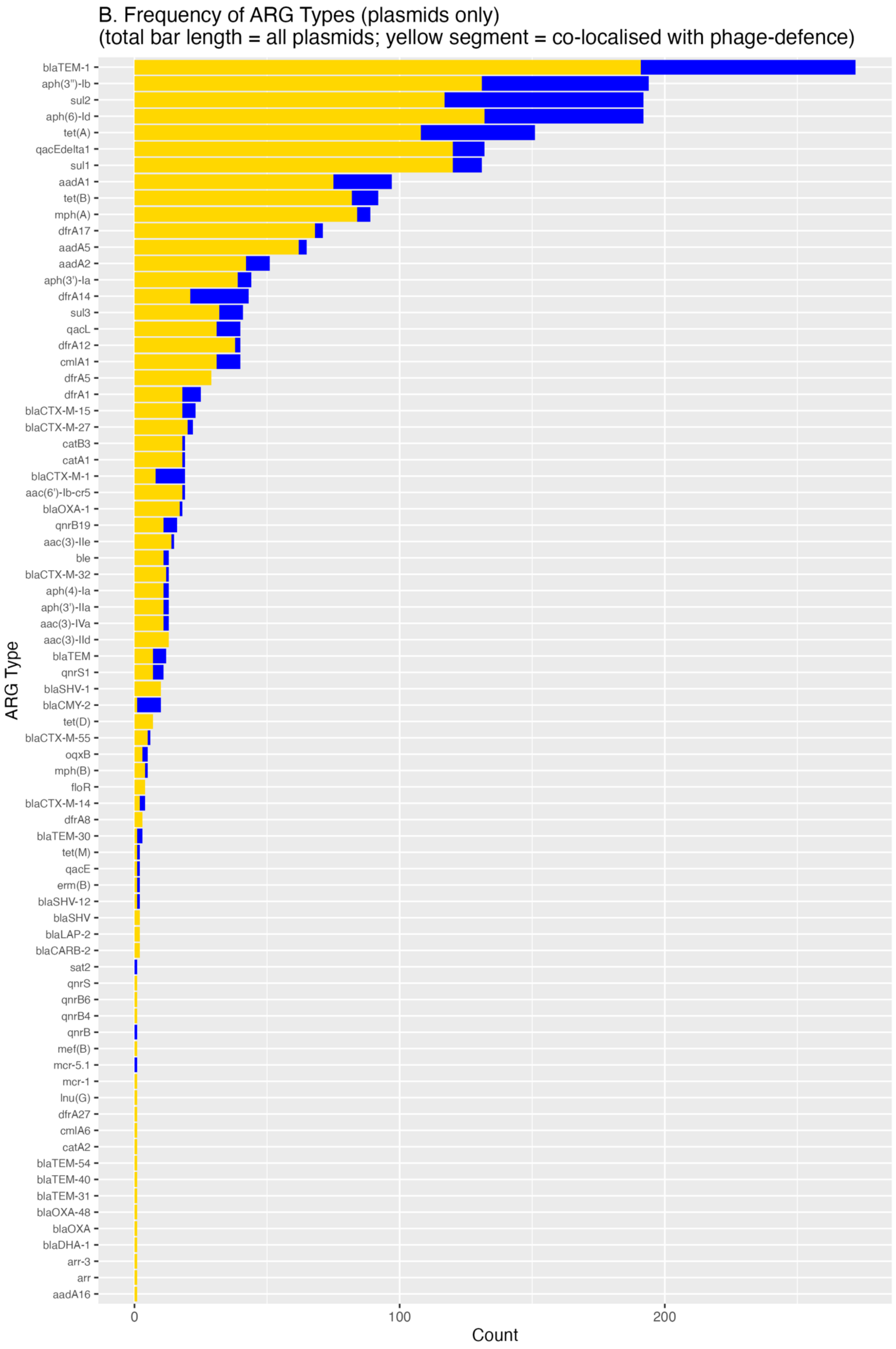
B. Frequency of ARG Types (plasmids only)
(total bar length = all plasmids; yellow segment = co-localised with phage-defence)
ARG Type
Count
0
100
200
blaTEM-1
aph(3'')-Ib
sul2
aph(6)-Id
tet(A)
qacEdelta1
sul1
aadA1
tet(B)
mph(A)
dfrA17
aadA5
aadA2
aph(3')-Ia
dfrA14
sul3
qacL
dfrA12
cmlA1
dfrA5
dfrA1
blaCTX-M-15
blaCTX-M-27
catB3
catA1
blaCTX-M-1
aac(6')-Ib-cr5
blaOXA-1
qnrB19
aac(3)-IIe
ble
blaCTX-M-32
aph(4)-Ia
aph(3')-IIa
aac(3)-IVa
aac(3)-IId
blaTEM
qnrS1
blaSHV-1
blaCMY-2
tet(D)
blaCTX-M-55
oqxB
mph(B)
floR
blaCTX-M-14
dfrA8
blaTEM-30
tet(M)
qacE
erm(B)
blaSHV-12
blaSHV
blaLAP-2
blaCARB-2
sat2
qnrS
qnrB6
qnrB4
qnrB
mef(B)
mcr-5.1
mcr-1
lnu(G)
dfrA27
cmlA6
catA2
blaTEM-54
blaTEM-40
blaTEM-31
blaOXA-48
blaOXA
blaDHA-1
arr-3
arr
aadA16

**Supplementary Figure S7. Relative frequencies of phage-defence systems and antimicrobial-resistance genes (ARGs) on plasmids.** For every phage-defence system type (panel A) and ARG type (panel B), horizontal bars give the number of plasmid contigs that carry at least one locus of that type. The total bar length (blue + yellow) is the count across all plasmids in the E. coli dataset, while the yellow segment is the subset of those plasmids on which the focal defence system and an ARG are co-localised on the same contig (panel A), or the focal ARG and a phage-defence system are co-localised (panel B). **(Panel A) Phage-defence systems.** The dataset contains 1,469 phage-defence loci on plasmids; 586 of them (40 %) occur on plasmids that also encode at least one ARG. The rank–frequency profile of the co-localised subset closely mirrors that of the full set: the most common systems (e.g. Mok_Hok_Sok, MazEF, RM) remain the most common after conditioning on ARG proximity, and the relative drop in counts is roughly uniform across the distribution. **(Panel B) ARGs.** Plasmids harbour 2,419 ARG loci in total, of which 1,864 (77 %) are found on contigs that also carry a phage-defence system. Although the abundance of individual ARG types declines when the analysis is restricted to co-localised contigs, their overall rank order is largely preserved (e.g. blaTEM-1, aph(3")-Ib, sul2, aph(6)-Id, tet(A), quacEdelta1, sul1, aadA1, tet(B), mph(A) remain the top ten). In both panels the strong visual concordance between the blue and yellow bar sequences indicates that co-localisation is probably not confined to a few specialist system types or ARGs, but broadly follows their background prevalence. Formal statistical testing of distributional differences was outside the scope of this study.

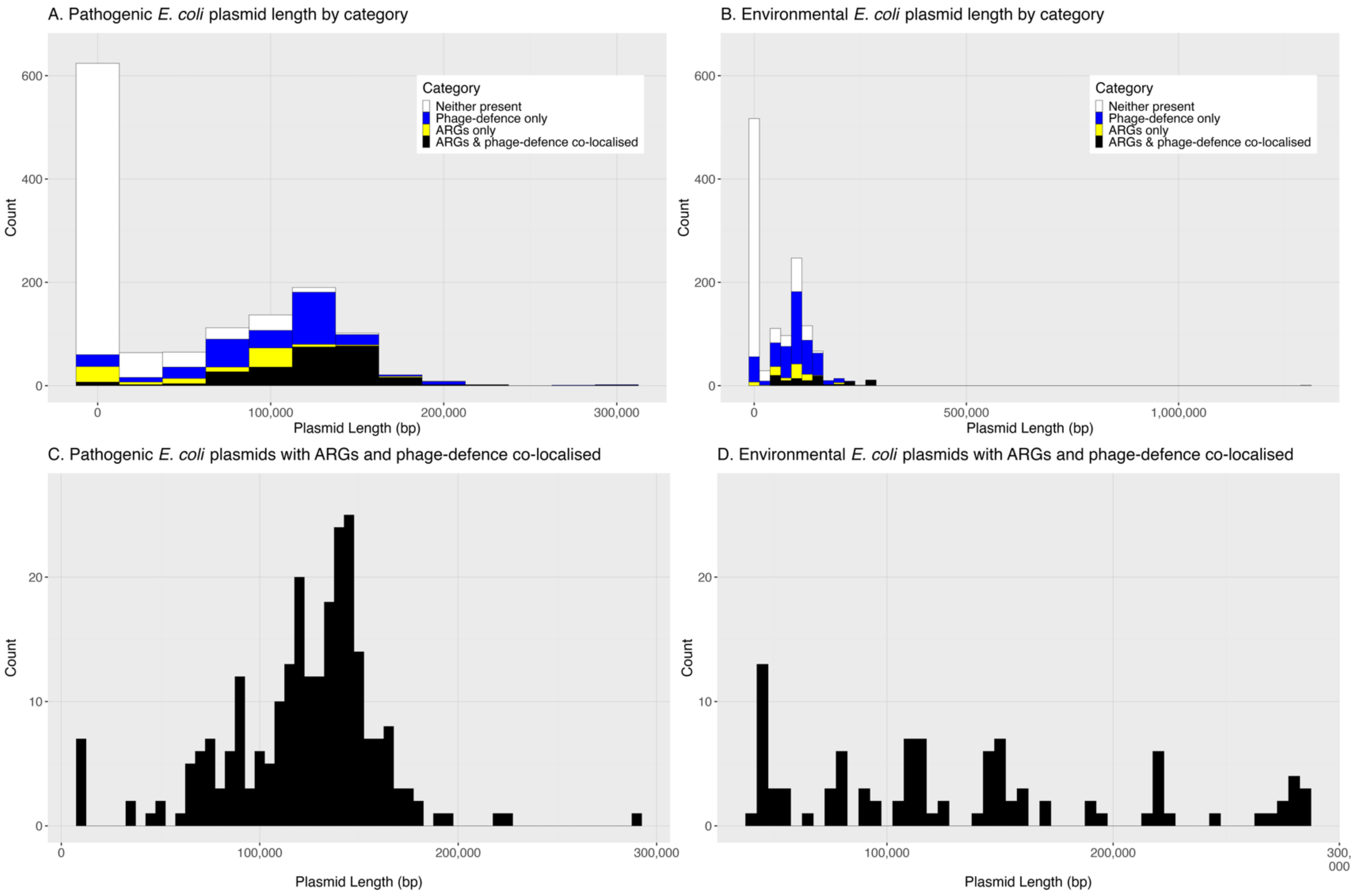


**Supplementary Figure S8**. **Length distribution of E. coli plasmids in relation to phage-defence systems and antibiotic-resistance genes (ARGs).** Panels show every plasmid contig in the study before sequence clustering; the common x-axis spans 0 – 300 kb (the largest plasmid observed) so that shapes can be compared directly across graphs. **(A) Pathogenic dataset (blood-stream-infection, BSI; n=1 328 plasmids).** Each 25 kb bin is split into four exclusive classes: plasmids that carry neither function (white), phage-defence only (blue), ARGs only (yellow) or plasmids that encode at least one ARG and one phage-defence system co-localised on the same contig (black). The y-axis is fixed at 0 – 650 on this and panel B to facilitate comparison. **(B) Environmental dataset (REHAB; n=1 229 plasmids)**. Plotted identically to panel A. The overall length distribution is more left-skewed than in the pathogenic collection, and almost all plasmids are < 150 kb despite the presence of a few very large environmental plasmids (> 1 Mb) at the extreme right tail. **(C) BSI plasmids in which ARGs and phage-defence systems are co-localised**. To reveal fine structure the bin width is reduced to 5 kb (60 bins across the 0 – 300 kb range) and the y-axis is rescaled to 0 – 27 (shared with panel D). The shape broadly mirrors panel A but retains a secondary peak of seven very small (< 13 kb) mobilisable ColRNAI_1 plasmids that harbour bla<sub>TEM-1</sub> together with RnlAB, AbiQ, Paris or RM defence modules. **(D) Environmental plasmids with co-localised ARGs and phage-defence systems**. Co-localisation is absent below ~44 kb in the environmental collection and the remaining plasmids display a multimodal distribution concentrated in the 45 - 200 kb range, indicating that linkage of the two functions in non-clinical reservoirs is restricted to medium- to large-sized plasmids. Across both sources the histograms emphasise that, while ARGs or phage-defence loci alone occur on plasmids of virtually any size, their co-localisation is strongly size-dependent—tolerated on the very smallest plasmids in pathogenic E. coli but restricted to substantially larger replicons in environmental isolates.

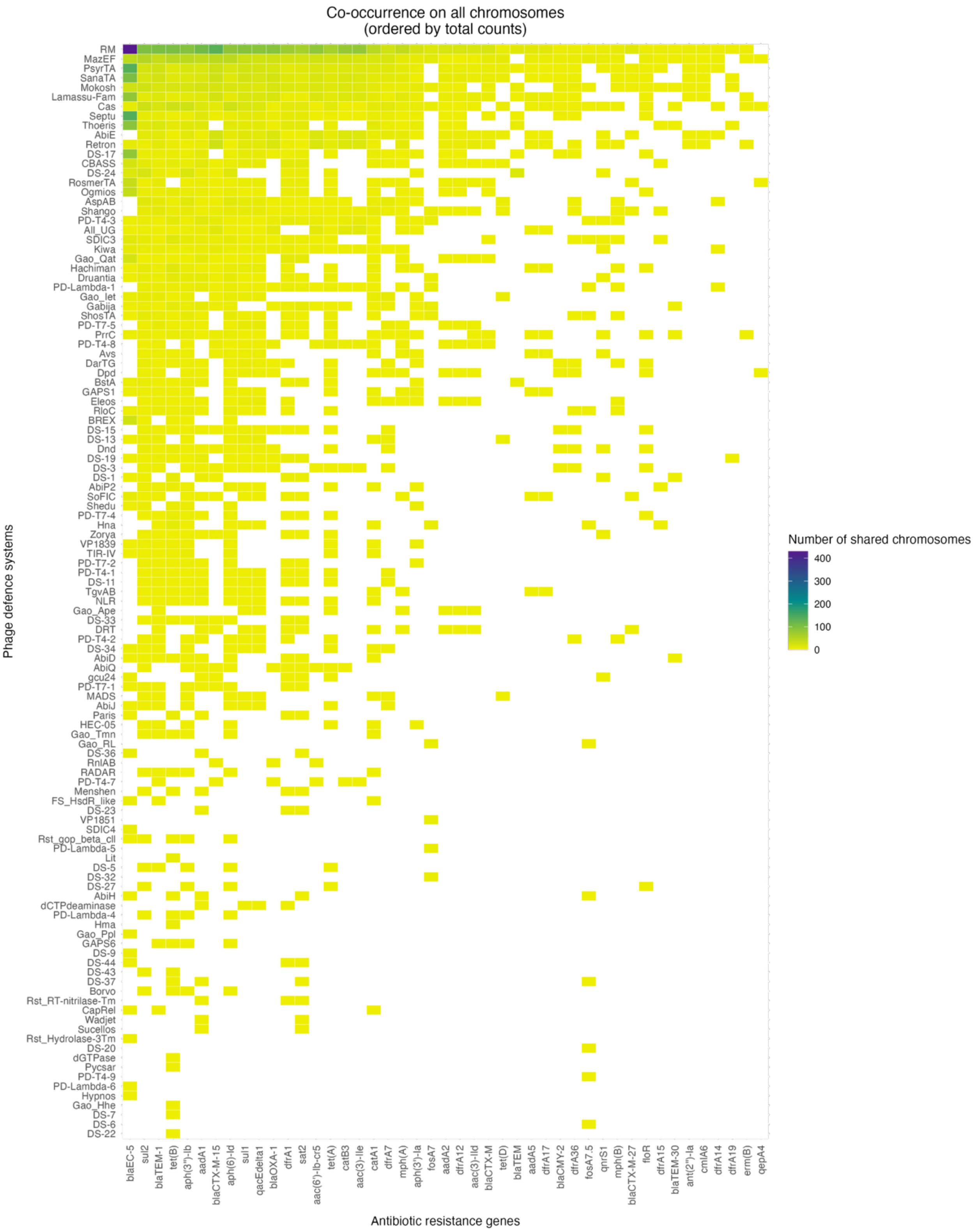


**Supplementary Figure S9. Heatmap presenting the frequency of co-localization of specific ARGs with specific phage-defence systems on the same chromosomes.** Each cell in the heatmap represents the count of chromosomes on which a given ARG (column) and phage-defence system (row) co-localize (the count of such chromosomes is indicated by the colour of the cell, according to the colour key in the upper left corner). ARG and phage-defence names are displayed as in DefenseFinder and AMRFinderPlus packages to aid in reproducibility. The figure presents the overall prevalence and not enrichment
