## Supplementary File 2 for "Plasmids link antibiotic resistance genes and phage defense systems in *E. coli*"

### GAMM modelling and Heatmap code

### Loading required packages

```
library(dplyr)
##
## Attaching package: 'dplyr'
## The following objects are masked from 'package:stats':
##
##     filter, lag
## The following objects are masked from 'package:base':
##
##     intersect, setdiff, setequal, union
library(mgcv)
## Warning: package 'mgcv' was built under R version 4.4.1
## Loading required package: nlme
##
## Attaching package: 'nlme'
## The following object is masked from 'package:dplyr':
##
##     collapse
## This is mgcv 1.9-3. For overview type 'help("mgcv-package")'.
library(broom)
## Warning: package 'broom' was built under R version 4.4.1
library(ggplot2)
## Warning: package 'ggplot2' was built under R version 4.4.1
library(scales)
library(ggtext)
library(tidyverse)
## Warning: package 'tidyr' was built under R version 4.4.1
## ── Attaching core tidyverse packages ──────────────────────── tidyverse 2.0.0 ──
## ✔ forcats   1.0.0     ✔ stringr   1.5.1
## ✔ lubridate 1.9.3     ✔ tibble    3.2.1
## ✔ purrr     1.0.2     ✔ tidyr     1.3.1
## ✔ readr     2.1.5
## ── Conflicts ────────────────────────────────────────── tidyverse_conflicts() ──
## ✖ readr::col_factor() masks scales::col_factor()
## ✖ nlme::collapse()    masks dplyr::collapse()
## ✖ purrr::discard()    masks scales::discard()
## ✖ dplyr::filter()     masks stats::filter()
## ✖ dplyr::lag()        masks stats::lag()
## ℹ Use the conflicted package (<http://conflicted.r-lib.org/>) to force all conflicts to become errors
library(rstatix)
##
## Attaching package: 'rstatix'
##
## The following object is masked from 'package:stats':
##
##     filter
library(DHARMa)
## This is DHARMa 0.4.6. For overview type '?DHARMa'. For recent changes, type news(package = 'DHARMa')
library(forcats)
library(gplots)
##
## Attaching package: 'gplots'
##
## The following object is masked from 'package:stats':
##
##     lowess
```

### Raw data processing

```
#Ensuring that PlasmidFinder NA values will also be included in the model

metadata_sampled_clusters <- metadata_sampled_clusters %>%
  mutate(
    Abricate.PlasmidFinder =
      fct_na_value_to_level(Abricate.PlasmidFinder, level = "Unknown")
  )
metadata_sampled_clusters <-
  metadata_sampled_clusters %>%
  mutate(
    # response & continuous predictors
    AMR_count               = as.integer(AMR_count),
    systems_count           = as.integer(systems_count),
    GC....                  = as.numeric(GC....),
    Length..bp.             = as.numeric(Length..bp.),
    # categorical predictors
    MOB.typer.mobility      = as.factor(MOB.typer.mobility),
    Niche                   = as.factor(Niche),
    Abricate.PlasmidFinder  = as.factor(Abricate.PlasmidFinder),
    REHAB.sample            = as.factor(REHAB.sample)
  )
metadata_sampled_clusters <-
  metadata_sampled_clusters %>%
  mutate(
    # numeric/continuous variables
    AMR_count      = as.integer(AMR_count),
    systems_count  = as.integer(systems_count),
    `GC....`       = as.numeric(`GC....`),
    `Length..bp.`  = as.numeric(`Length..bp.`),

    # categorical variables
    MOB.typer.mobility     = as.factor(MOB.typer.mobility),
    Niche                  = factor(Niche,
                                     levels = c("Livestock-associated",
                                                "BSI",
                                                "WwTW-associated")),
    Abricate.PlasmidFinder = as.factor(Abricate.PlasmidFinder)
  ) %>%

  # centre GC so that 0 = mean(GC)
  mutate(
    GC_mean      = mean(`GC....`, na.rm = TRUE),     # save the mean (for reporting)
    GC_centered  = `GC....` - GC_mean                # mean-centred version used in the model
  )
```

### GAMM fitting

```
# Generalised additive model (Tweedie, log link)
gamm_model <- gam(
  AMR_count ~
    systems_count +
    GC_centered +                                # centred GC predictor
    offset(log10(`Length..bp.`)) +
    Niche +                                      # reference = "Livestock-associated"
    s(Abricate.PlasmidFinder, bs = "re"),
  family = tw(link = "log"),
  data   = metadata_sampled_clusters
)

# Model inspection
summary(gamm_model)
##
## Family: Tweedie(p=1.289)
## Link function: log
##
## Formula:
## AMR_count ~ systems_count + GC_centered + offset(log10(Length..bp.)) +
##     Niche + s(Abricate.PlasmidFinder, bs = "re")
##
## Parametric coefficients:
##                      Estimate Std. Error t value Pr(>|t|)
## (Intercept)          -6.58008    0.34009 -19.348  < 2e-16 ***
## systems_count         0.39108    0.11678   3.349 0.000856 ***
## GC_centered           0.31802    0.03785   8.403 2.54e-16 ***
## NicheBSI              0.72005    0.26960   2.671 0.007748 **
## NicheWwTW-associated  0.47746    0.46328   1.031 0.303090
## ---
## Signif. codes:  0 '***' 0.001 '**' 0.01 '*' 0.05 '.' 0.1 ' ' 1
##
## Approximate significance of smooth terms:
##                            edf Ref.df    F p-value
## s(Abricate.PlasmidFinder) 25.7     57 1.21 5.9e-07 ***
## ---
## Signif. codes:  0 '***' 0.001 '**' 0.01 '*' 0.05 '.' 0.1 ' ' 1
##
## R-sq.(adj) =  0.402   Deviance explained = 42.1%
## -REML = 410.76  Scale est. = 4.0278    n = 712
AIC(gamm_model)
## [1] 810.8677
# DHARMa for Tweedie
sim_twp <- DHARMa::simulateResiduals(gamm_model)
## Registered S3 method overwritten by 'GGally':
##   method from
##   +.gg   ggplot2
## Registered S3 method overwritten by 'mgcViz':
##   method from
##   +.gg   GGally
DHARMa::plotSimulatedResiduals(sim_twp)
## plotSimulatedResiduals is deprecated, please switch your code to simply using the plot() function
```

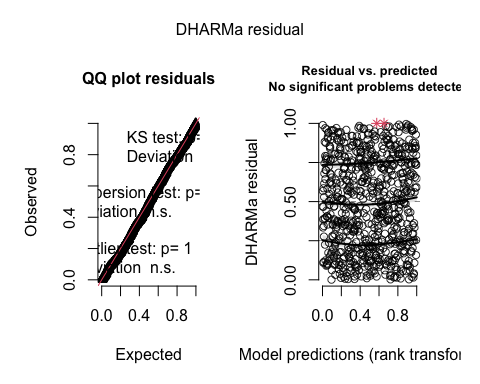

DHARMa::testDispersion(sim_twp)

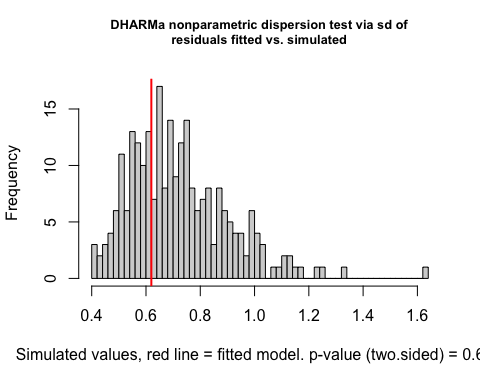

```
##
##  DHARMa nonparametric dispersion test via sd of residuals fitted vs.
##  simulated
##
## data:  simulationOutput
## dispersion = 0.8584, p-value = 0.656
## alternative hypothesis: two.sided
```

### Forest plot visualisation

```
# Forest-plot of gamm_model

# 1. Tidy coefficients & exponentiate (to fold-changes)
coef_df <- tidy(gamm_model, parametric = TRUE, conf.int = TRUE) %>%
  filter(term != "(Intercept)") %>%
  mutate(across(c(estimate, conf.low, conf.high), exp))   # exp

# 2. Add the “reference” row (fold-change = 1)
ref_lvl <- levels(model.frame(gamm_model)$Niche)[1]            # "Livestock-associated"
coef_df <- bind_rows(
  tibble(
    term      = paste0("Niche", ref_lvl, " (reference)"),
    estimate  = 1,
    conf.low  = 1,
    conf.high = 1
  ),
  coef_df
)

#  3. Human-readable labels & desired plotting order
coef_df <- coef_df %>%
  mutate(
    label = case_when(
      term == "systems_count"                     ~ "Phage-defense systems count",
      term == "GC_centered"                       ~ "GC content (mean-centred)",
      term == paste0("Niche", ref_lvl, " (reference)")
                                                 ~ "Niche: Livestock-associated (reference)",
      term == "NicheWwTW-associated"              ~ "Niche: Wastewater",
      term == "NicheBSI"                          ~ "Niche: Bloodstream infections",
      TRUE                                        ~ term
    )
  )

desired_order <- c(
  "Phage-defense systems count",
  "GC content (mean-centred)",
  "Niche: Livestock-associated (reference)",
  "Niche: Wastewater",
  "Niche: Bloodstream infections"
)

coef_df <- coef_df %>%
  mutate(label = factor(label, levels = rev(desired_order)))

# 4. Highlight the phage-defense row & build markdown labels
coef_df <- coef_df %>%
  mutate(highlight = (label == "Phage-defense systems count"))

lab_vec <- levels(coef_df$label)
lab_out <- ifelse(
  lab_vec == "Phage-defense systems count",
  "<span style='color:blue2;font-weight:bold'>Phage-defense systems count</span>",
  lab_vec
)

# 5. Build the plot (linear x-axis)
pA <- ggplot(coef_df, aes(x = estimate, y = label, colour = highlight)) +
  geom_vline(xintercept = 1, linetype = "dashed") +
  geom_errorbarh(aes(xmin = conf.low, xmax = conf.high), height = 0.2) +
  geom_point(size = 3) +
  scale_colour_manual(values = c(`TRUE` = "blue2", `FALSE` = "black"), guide = "none") +
  scale_x_continuous(
    breaks = c(0.5, 1, 1.5, 2, 4, 8),            # linear axis with extra ticks
    labels = number_format(accuracy = 0.1)
  ) +
  scale_y_discrete(labels = lab_out) +
  labs(
    title = "A. Generalised additive model predictor effects on ARG count\n(fold-change)",
    x     = "Fold-change (95 % CI) in ARG count",
    y     = NULL
  ) +
  theme_classic(base_size = 14) +
  theme(
    panel.grid.major   = element_line(colour = "grey85", size = 0.2),
    panel.grid.minor.x = element_line(colour = "grey90", size = 0.1),
    axis.text.x  = element_text(size = 13),
    axis.text.y  = element_markdown(size = 13),
    axis.title.x = element_text(size = 15, face = "bold"),
    axis.title.y = element_text(size = 15, face = "bold"),
    plot.title   = element_text(size = 16, face = "bold", hjust = 0.5)
  )
## Warning: The `size` argument of `element_line()` is deprecated as of ggplot2 3.4.0.
## ℹ Please use the `linewidth` argument instead.
## This warning is displayed once every 8 hours.
## Call `lifecycle::last_lifecycle_warnings()` to see where this warning was
## generated.
# 6. Also print to the current graphics device (optional)
pA
```

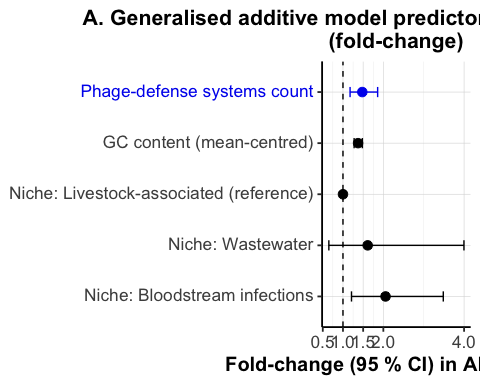

### Multicollinearity checks

```
## 1. Extract the data actually used in the fit
df <- model.frame(gamm_model)

## 2. Identify variables
# Anything that begins with "offset(" is dropped
# Any variable that is the random effect is dropped too (here: Abricate.PlasmidFinder)
offset_vars   <- names(df)[grepl("^offset\\(", names(df))]
random_effect <- "Abricate.PlasmidFinder"

cont_vars <- names(df)[map_lgl(df, is.numeric)] |> setdiff(offset_vars)
cat_vars  <- names(df)[map_lgl(df, ~ is.factor(.x) || is.character(.x))] |>
             setdiff(random_effect)

## 3. Continuous × continuous  (Spearman ≥ 0.7)
spearman_mat <- df |>
  select(all_of(cont_vars)) |>
  cor(method = "spearman", use = "pairwise.complete.obs")

spearman_flags <- as.data.frame(as.table(spearman_mat)) |>
  mutate(across(c(Var1, Var2), as.character)) |>      # avoid factor warning
  filter(Var1 < Var2) |>
  mutate(abs_rho = abs(Freq), flag = abs_rho >= 0.7) |>
  arrange(desc(abs_rho))

cat("\n--- Continuous × Continuous ( ≥ 0.7) ---\n")
##
## --- Continuous × Continuous ( ≥ 0.7) ---
print(spearman_flags |> filter(flag) |>
        select(var1 = Var1, var2 = Var2, rho = Freq, abs_rho))
## [1] var1    var2    rho     abs_rho
## <0 rows> (or 0-length row.names)
## 4. Continuous × categorical (Kruskal–Wallis ≥ 0.7)
kw_results <- expand_grid(cont = cont_vars, cat = cat_vars) |>
  mutate(
    kw_test = map2(cont, cat, ~ kruskal.test(df[[.x]], df[[.y]])),
    H       = map_dbl(kw_test, "statistic"),
    N       = map_dbl(cont,   ~ sum(!is.na(df[[.x]]))),
    eta_sq  = H / (N - 1),
    flag    = eta_sq >= 0.7
  ) |>
  arrange(desc(eta_sq))

cat("\n--- Continuous × Categorical ( ≥ 0.7) ---\n")
##
## --- Continuous × Categorical ( ≥ 0.7) ---
print(kw_results |> filter(flag) |> select(cont, cat, eta_sq))
## # A tibble: 0 × 3
## # ℹ 3 variables: cont <chr>, cat <chr>, eta_sq <dbl>
```

### Heatmap

```
# Keep only plasmid entries
plasmids <- meta %>%
  filter(Type == "Plasmid")

# Split the AMRFinderPlus column on commas
plasmids_long <- plasmids %>%
  mutate(AMRFinderPlus = coalesce(AMRFinderPlus, "")) %>%
  separate_rows(AMRFinderPlus, sep = ",") %>%
  mutate(AMRFinderPlus = str_trim(AMRFinderPlus)) %>%
  filter(AMRFinderPlus != "", !is.na(type))

# Build the co-occurrence matrix (ARG × defence)
co_mat <- table(plasmids_long$AMRFinderPlus, plasmids_long$type)

# Remove genes/systems that never co-occur
co_mat <- co_mat[rowSums(co_mat) > 0, colSums(co_mat) > 0]

# Convert to long format and compute totals
co_df <- as.data.frame(as.table(co_mat), stringsAsFactors = FALSE)
colnames(co_df) <- c("gene", "system", "count")

gene_totals   <- co_df %>% group_by(gene)   %>% summarise(total = sum(count))
system_totals <- co_df %>% group_by(system) %>% summarise(total = sum(count))

gene_levels   <- gene_totals   %>% arrange(desc(total)) %>% pull(gene)
system_levels <- system_totals %>% arrange(total)        %>% pull(system)

co_df <- co_df %>%
  mutate(
    gene   = factor(gene,   levels = gene_levels),
    system = factor(system, levels = system_levels)
  )

# Build the same colour palette as in your original heatmap.2() call
pal <- c("white", colorpanel(10000, "yellow2", "darkcyan", "purple4"))

# Draw the heatmap with ggplot2
pB <- ggplot(co_df, aes(x = gene, y = system, fill = count)) +
  geom_tile(color = "white") +
  scale_fill_gradientn(
    colours = pal,
    name    = "Number of shared plasmids"
  ) +
  labs(
    x     = "Antibiotic resistance genes",
    y     = "Phage defense systems",
    title = "B. Co-occurrence on all plasmids\n(ordered by total counts)"
  ) +
  theme_minimal(base_size = 14) +
  theme(
    plot.title    = element_text(size = 16, face = "bold", hjust = 0.5),
    axis.text.x   = element_text(angle = 90, vjust = 0.5, hjust = 1),
    axis.text.y   = element_text(hjust = 1),
    axis.title.y  = element_text(margin = margin(r = 0), face = "bold"),
    axis.title.x  = element_text(margin = margin(t = 10), face = "bold"),
    legend.position = "bottom"
  )

print(pB)
```

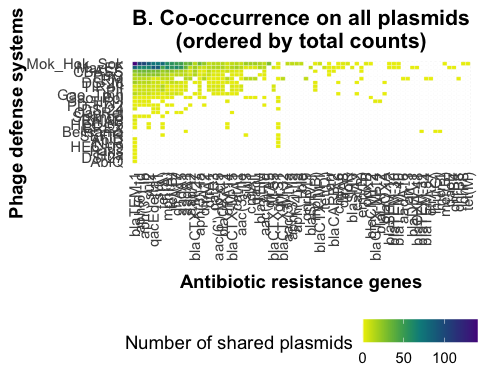
